# SHINE: Decoding transcriptional-metabolic microenvironments through higher-order spatial integration

**DOI:** 10.64898/2026.07.10.737648

**Authors:** Bingxue Du, Jason Wing Hon Wong, Yuanhua Huang

## Abstract

Spatial omics technologies are expanding to co-profile transcriptomics and metabolomics on the same tissue slide, providing complementary views of gene expression and biochemical activity to reveal molecular programs within native tissue microenvironments. However, integrating the transcriptome and metabolome remains technically challenging due to spatial misalignment, resolution disparity, and higher-order cross-modality interactions. Here, we present **SHINE**, a hypergraph-based computational framework for the joint analysis of spatial gene expression and metabolic networks derived from the co-profiling slide, focusing on representation learning and cross-modality interaction. Across multiple datasets, SHINE consistently outperformed existing methods for domain segmentation and biomarker co-localization, and provided interpretable insights into metabolic–transcriptional microenvironments. Specifically, on Parkinson’s disease mouse models, SHINE accurately delineates dopaminergic neuron-depleted regions and reconstructs coherent dopamine-associated axes. In human lung and breast cancers, SHINE resolves tumor-associated spatial regions and identifies spatially organized gene–metabolite programs associated with the tumor microenvironment. SHINE enables scalable spatial multi-omics integration across diverse biological systems.

## 1 Main

Spatial organization is fundamental to tissue biology, shaping how molecular programs and cell states are organized within native microenvironments to drive physiological function and disease [1–3]. Recent advances in spatial omics technologies now enable in situ profiling of tissues with multi-cellular to sub-cellular resolution [4]. Spatial transcriptomics (ST) captures gene expression landscapes and can further reveal transcript isoform diversity arising from alternative splicing [5], whereas spatial metabolomics (SM), such as mass spectrometry imaging (MSI), directly measures the spatial distribution of metabolites reflecting cellular biochemical activity [6]. Together, these complementary modalities provide a powerful foundation for dissecting complex tissue phenotypes [7].

Integrating SM and ST has begun to reveal disease-associated molecular programs, including dopaminergic metabolic alterations in Parkinson’s disease (PD) [8] and metabolic reprogramming within tumor microenvironments [9, 10]. However, computational integration remains challenging, including spatial-resolution mismatch, imperfect cross-modality registration, and batch effects across samples, which collectively hinder coherent modeling of coordinated molecular processes within tissue microenvironments [11]. Recent advances have also explored histology-guided strategies to facilitate spatial multi-omics integration across tissue sections [12]. Most existing methods rely on feature concatenation, latent alignment, or first-order (pairwise) spatial graphs, which implicitly reduce tissue organization to collections of one-to-one spot-pair relationships.

Several computational frameworks have been proposed to address spatial multi-modality integration. Graph-based methods such as SpatialGlue [13] leverage spatial adjacency and feature similarity to align modalities, whereas recent approaches, including MISO [14] and SpatialMETA [15], employ latent variable and variational modeling strategies to integrate spatial metabolomics and transcriptomics across samples. While these methods effectively capture local correspondence between modalities, they primarily operate at the level of individual spots or spot pairs, modeling cross-modality relationships independently within local neighborhoods. In biological tissues, however, functional microenvironments are not defined by isolated spatial units [16], but instead emerge from the coordinated activity of gene programs and metabolic pathways across groups of spatially proximal spots (e.g., niches or domains) [17]. Such a region-level organization reflects collective, many-to-many interactions that cannot be decomposed into independent spot-pairwise relationships. These considerations motivate higher-order interaction modeling frameworks, such as hypergraph learning, which can naturally encode collective spatial relationships and better align computational modeling with the organization principles of tissue microenvironments [18].

To address these challenges, we introduce SHINE (Spatial Hypergraph Integration of metabolic Networks and gene Expression), a hypergraph-based framework for integrative analysis of spatial metabolomics and transcriptomics. SHINE represents coordinated RNA-metabolite programs and their spatial context as higher-order relational structures, enabling principled modeling of tissue microenvironments beyond pairwise spot-level associations. The framework explicitly captures multi-way metabolic–transcriptional interactions embedded within spatially organized tissues by integrating brightfield-guided cross-modality registration, modality-specific spatial and feature graphs, graph transformer layers for long-range dependency modeling, and a causally masked cross-modality hypergraph. Together, these components learn coherent higher-order multi-modal embeddings that reflect region-level molecular coordination.

We applied SHINE to multiple datasets with paired spatial profiling of RNA-metabolite spanning species, tissues, and disease contexts, including mouse striatum and substantia nigra (SN) in unilateral 6-OHDA PD models, as well as human lung cancer and breast cancer. Using the default parameter configuration across all datasets, SHINE consistently outperformed state-of-the-art methods in recovering spatial architecture and biologically coherent domains. SHINE delineated lesion-associated microenvironments in PD and resolved tumor–stroma organization and metabolic programs in human cancers. In summary, SHINE addresses a central limitation of current spatial multi-omics methods by explicitly modeling higher-order multi-modality interactions beyond pairwise representations.

## 2 Results

### 2.1 SHINE: a hypergraph integration framework for spatial metabolomics and transcriptomics

We present SHINE (Fig. 1), a spatial omics framework that enables cross-modality metabolic–transcriptional integration and microenvironment reconstruction across tissues and species (Fig. 1A). Tissue sections from mouse and human samples are profiled with MSI for metabolite abundance and Visium for gene expression. We aligned MSI pixels with Visium spots using brightfield-guided rigid and thin-plate spline (TPS) transformations, thereby generating a unified spatial framework that enables metabolite–gene pairing at the Visium spot level.

**Fig. 1.**
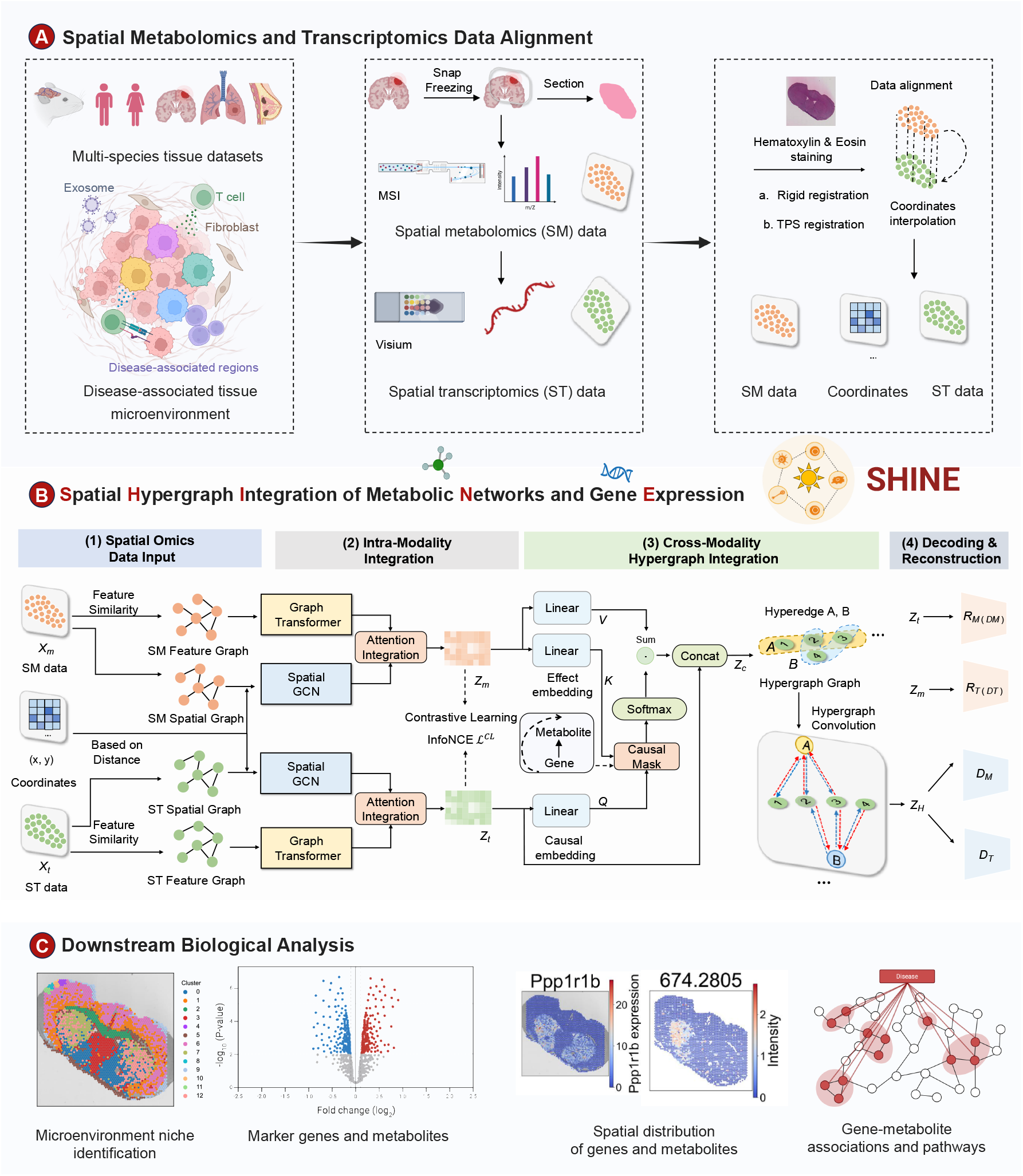
Overview of SHINE for cross-modality spatial metabolomics and transcriptomics integration. **A**, Spatial Metabolomics and Transcriptomics Data Alignment. Tissue sections spanning multiple species, organs and disease contexts are profiled using SM and ST. Histology-guided rigid and TPS registration aligns metabolite pixels and transcriptomic spots into a unified spatial coordinate system, generating paired metabolite–gene profiles across tissue locations and enabling characterization of spatial disease microenvironments. **B**, Spatial Hypergraph Integration of Metabolic Networks and Gene Expression. **(1)**, Spatial Omics Data Input. The aligned SM and ST matrices are converted into modality-specific spatial graphs (based on spot coordinates) and feature graphs (based on metabolite intensities or gene expression similarity); **(2)**, Intra-Modality Integration. SM and ST modalities are independently encoded using spatial GCNs and graph Transformer layers, generating modality-specific embeddings that capture both local spatial structure and long-range tissue organization; **(3)** Cross-Modality Hypergraph Integration. A cross-modality hypergraph is constructed using KNN-based feature similarity and causal masking to link groups of spatially or functionally coherent spots across modalities. Hypergraph convolution produces an integrated embedding that models higher-order gene–metabolite interactions and microenvironment-level regulation. **(4)** Decoding & Reconstruction. Modality-specific GCN decoders generate ST and SM predictions, while Graph Transformer modules reconstruct modality-specific features. **C**, Downstream Biological Analyses. Created in part with BioRender.com.

Based on the aligned RNA-metabolite input, SHINE converts SM and ST data into modality-specific spatial and feature graphs (Fig. 1B). The framework comprises four sequential components. First, paired SM and ST datasets are co-registered and curated to ensure spatial correspondence across modalities. Second, modality-specific spatial-proximity graphs and feature-similarity graphs are constructed for SM and ST. Each modality is independently encoded using a spatial Graph Transformer and SpatialGCN layers to capture feature similarity and spatial proximity. Third, cross-modality relationships are integrated through a hypergraph-based formulation, in which groups of spatially proximal or functionally coherent spots are linked across modalities using *k*-nearest neighbor (KNN)–based feature similarity and causal masking, thereby enabling explicit modeling of higher-order gene–metabolite interactions. Finally, modality-specific decoders reconstruct SM and ST features as well as spatial representations, and a composite objective jointly optimizes within-modality reconstruction fidelity and cross-modality consistency for downstream inference of spatially resolved tissue microenvironments. The resulting integrated embeddings enable downstream analyses such as spatial microenvironment niche identification, marker gene and metabolite discovery, and inference of gene–metabolite associations and pathways, facilitating biological interpretation of spatially resolved tissue microenvironments (Fig. 1C). Additional methodological details are provided in the Methods and Supplementary Information.

To assess robustness and generalizability, we benchmarked SHINE across diverse spatial RNA-metabolite settings, spanning two species, three disease contexts, four tissue types, and five spatial slides. All methods were implemented following their recommended parameter settings. Performance was quantified against region annotations or cell-type labels using Adjusted Rand Index (ARI) and Normalized Mutual Information (NMI), in comparison with three recently proposed methods: SpatialGlue [13], MISO [14], and SpatialMETA [15].

To evaluate the effectiveness of the proposed SHINE framework, we first examined the impact of multi-omics integration on spatial and UMAP-based clustering results by comparing single-modality and integrated representations (Supplementary Fig. 1), which consistently produced clearer cluster structure and improved agreement with biological annotations. We then assessed the contribution of hypergraph-based integration and KNN-based fusion through a series of ablation studies (Supplementary Fig. 2), which indicated that hypergraph-based integration enables more fine-grained modeling of higher-order relationships and leads to improved clustering performance. Having established its robustness and modeling advantages, we next applied SHINE to representative disease settings, including Parkinson’s disease, lung cancer, and breast cancer, to examine how integrative spatial modeling resolves disease-associated molecular organization.

### 2.2 SHINE resolves spatial metabolic–transcriptional alterations in the mouse striatum of a Parkinson’s disease model

To evaluate how well SHINE captures metabolic–transcriptional alterations associated with neurodegeneration, we applied it to a unilateral 6-hydroxydopamine (6-OHDA) lesioned mouse striatum—a widely used PD model that causes selective dopaminergic neuron loss [19]. SM measurements were acquired using matrix-assisted laser desorption/ionization mass spectrometry imaging (MALDI–MSI) with FMP-10 as the matrix. Using anatomical region annotations from the original study [8] and its single-cell derived cell-type labels as ground truth (Fig. 2a), SHINE accurately delineated PD-affected territories, including the dopamine-depleted medial striatum. In contrast, baseline multi-omics integration methods [13–15] produced fragmented or anatomically inconsistent domain boundaries (Fig. 2b). Quantitative comparisons confirmed that SHINE achieved the highest ARI and NMI scores across region labels, cell types, and PD-relevant categories, with a substantial gain, for example, ARI for the anatomical region (0.423 by SHINE vs 0.161 by the best counterpart; Fig. 2c).

**Fig. 2.**
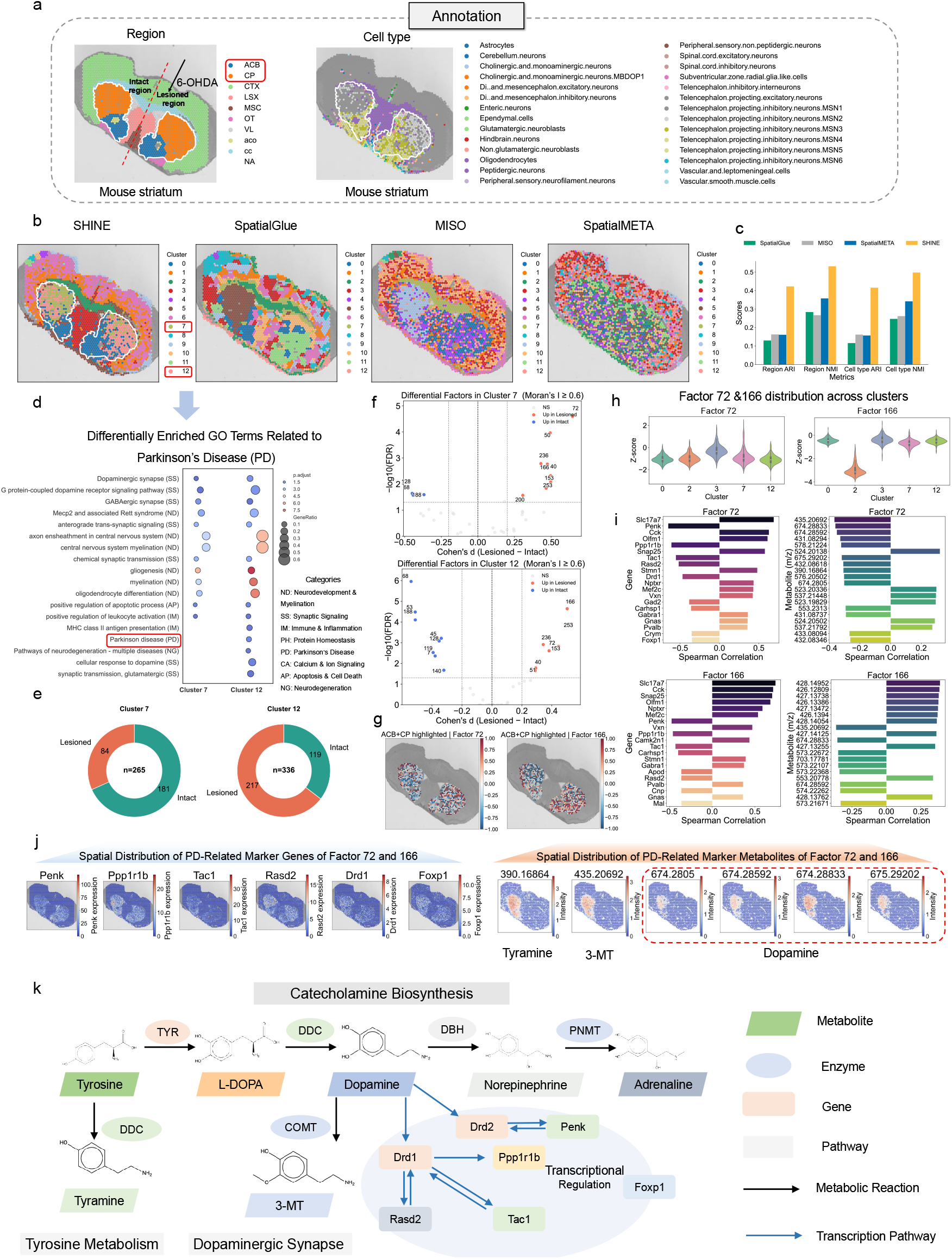
Comparative spatial multi-omics analysis of the PD mouse striatum using SHINE. **A**, Anatomical region annotations and cell-type labels of the mouse striatum following unilateral 6-OHDA lesioning. **b**, Spatial clustering results generated by SHINE, SpatialGlue, MISO and SpatialMETA. **c**, Quantitative evaluation of clustering performance using ARI and NMI with respect to anatomical regions, major cell types and PD-related annotations. **d**, GO terms enriched among genes associated with PD-related spatial domains identified by SHINE. **e**, Proportions of spatial clusters enriched in the lesioned versus contralateral hemispheres. **f**, Differential analysis of latent factors in PD-associated clusters (Clusters 7 and 12), showing effect sizes (Cohen’s d, lesioned versus intact hemisphere) and statistical significance for factors with strong spatial autocorrelation (Moran’s *I ≥* 0.6). **g**, Latent metabolic–transcriptional factors (Factors 72 and 166) learned by SHINE, shown with their spatial distributions and ranked gene and metabolite loadings. **h**, Distribution of Factors 72 and 166 across spatial clusters. **i**, Top gene and metabolite loadings for Factors 72 and 166. **j**, Spatial distribution of PD-related marker genes and metabolites associated with Factors 72 and 166. **k**, Reconstructed catecholamine biosynthesis and dopaminergic synapse pathways integrating SHINE-derived gene–metabolite associations (KEGG module: mmu M00042).

We next explored whether the molecular features identified by SHINE correspond to known PD-related biology. By focusing on the caudoputamen (CP) region and its overlapped Clusters 7 and 12, we identified a set of differentially expressed genes between clusters, followed by the gene ontology (GO) enrichment analysis. These results revealed that genes upregulated in Cluster 7 were selectively associated with pathways relevant to dopaminergic signaling, neuroinflammation, and synaptic remodeling, whereas Cluster 12 lacked such associations (Fig. 2d). Spatial distribution across hemispheres confirmed that Cluster 7 was highly lesion-enriched (Fig. 2e). These cluster-driven features were further supported by distinct sets of marker genes and metabolites that characterized Clusters 7 and 12, respectively (Supplementary Fig. 3), highlighting their divergent molecular composition.

Latent factor analysis further uncovered PD-associated axes that mapped onto these lesion-specific domains. These latent factors, corresponding to dimensions of the integrative latent embedding (*Z*_*H*_) that capture coordinated spatial gene-metabolite programs, revealed structured molecular organization within lesion-associated domains. Among them, Factor 72 and Factor 166 stood out for their strong spatial localization and differential loadings between hemispheres (Fig. 2f). Differential analysis further identified a subset of spatially coherent latent factors that were significantly enriched in either the lesioned or intact hemisphere within PD-associated clusters. Notably, both factors were concentrated within the dopamine-depleted medial striatum, as shown by their spatial distributions (Fig. 2g). GO enrichment analysis of highly weighted genes in Factor 72 and 166 revealed strong associations with PD-relevant biological processes, including dopamine metabolism, synaptic signaling, and neuroinflammation (Supplementary Fig. 4), further supporting their mechanistic relevance to PD pathology. At the cluster level, these factors displayed selective enrichment in PD-associated spatial clusters, with distinct activation patterns across clusters (Fig. 2h), supporting their association with lesion-specific microenvironments. Their top gene and metabolite loadings (Fig. 2i) revealed coordinated multiple patterns in which neuropeptide expression, dopamine signaling genes, and related metabolic features showed aligned loading profiles.

To better interpret the latent factor–associated molecular programs, we visualized top-ranked genes and metabolites correlated with PD-enriched latent factors such as Factor 72 and Factor 166. These features exhibited coherent spatial localization patterns centered on the lesioned striatum (Supplementary Fig. 5). Notably, several gene–metabolite pairs—including *Penk, Ppp1r1b, Tac1, Rasd2, Drd1, Foxp1*, and metabolites (annotations listed in Supplementary Table S1) showed pronounced lateral asymmetry across hemispheres, further supporting lesion-specific co-enrichment as observed in Fig. 2j. These jointly localized signatures reinforce the biological relevance of SHINE-derived domains. Many of these genes are known mediators of dopaminergic signaling and striatal circuit remodeling in PD. For instance, *Ppp1r1b* and *Drd1* are critical for dopaminoceptive control of basal ganglia output, while *Penk* and *Tac1* reflect neuropeptide plasticity following dopamine depletion. *Rasd2*, encoding Rhes, is selectively enriched in striatal neurons, and its downregulation is a conserved response to dopamine loss in PD models.

To further validate the biological interpretability of these features, we integrated gene–metabolite associations into biochemical pathways. SHINE reconstructed a unified view of catecholamine biosynthesis and dopaminergic synapse organization (KEGG: mmu M00042), highlighting convergent axes centered on *Penk, Ppp1r1b, Tac1, Rasd2, Drd1*, and *Foxp1* (Fig. 2k). These results demonstrate that SHINE effectively unifies multi-modal features into coherent pathways aligned with PD pathophysiology.

To assess generalizability across experimental conditions, we further applied SHINE to an independent spatial metabolomics dataset generated from the same tissue section using another MALDI matrix, 2,5-dihydroxybenzoic acid (DHB). The method maintained consistent spatial clustering and domain delineation, supporting robustness across sample preparation and ionization protocols (Supplementary Fig. 6). SHINE also consistently outperformed baseline integration methods under the DHB matrix setting.

In summary, SHINE delineates lesion-specific multi-modal domains in the striatum and resolves spatial metabolic–transcriptional programs consistent with established PD mechanisms. Together, our findings demonstrate SHINE’s capacity to decode biologically meaningful microenvironments underlying Parkinsonian pathology. Given that Parkinson’s pathology involves coordinated degeneration along the nigrostriatal axis, we next asked whether SHINE could resolve analogous spatial metabolic–transcriptional programs in the SN, the primary site of dopaminergic neuron vulnerability.

### 2.3 Cross-regional mapping of Parkinson’s disease molecular programs in the substantia nigra by SHINE

We next applied SHINE to the SN, another brain region critically affected in PD, using the unilateral 6-OHDA lesion mouse model (Fig. 3). Using region annotations and cell-type labels for the lesioned vs. intact hemispheres (Fig. 3a) as references, we found that SHINE again produced coherent, anatomically interpretable clusters. In particular, it accurately delineated the dopaminergic neuron–depleted SN pars compacta (SNc), which is the primary site of neuron loss in PD. Baseline methods failed to capture this structure, yielding fragmented or mis-merged domains that did not align with known anatomy (Fig. 3b). Quantitatively, SHINE achieved superior ARI/NMI scores compared to all existing methods across region labels, major cell types and PD-related categories (Fig. 3c), underscoring its robust performance across tissues.

**Fig. 3.**
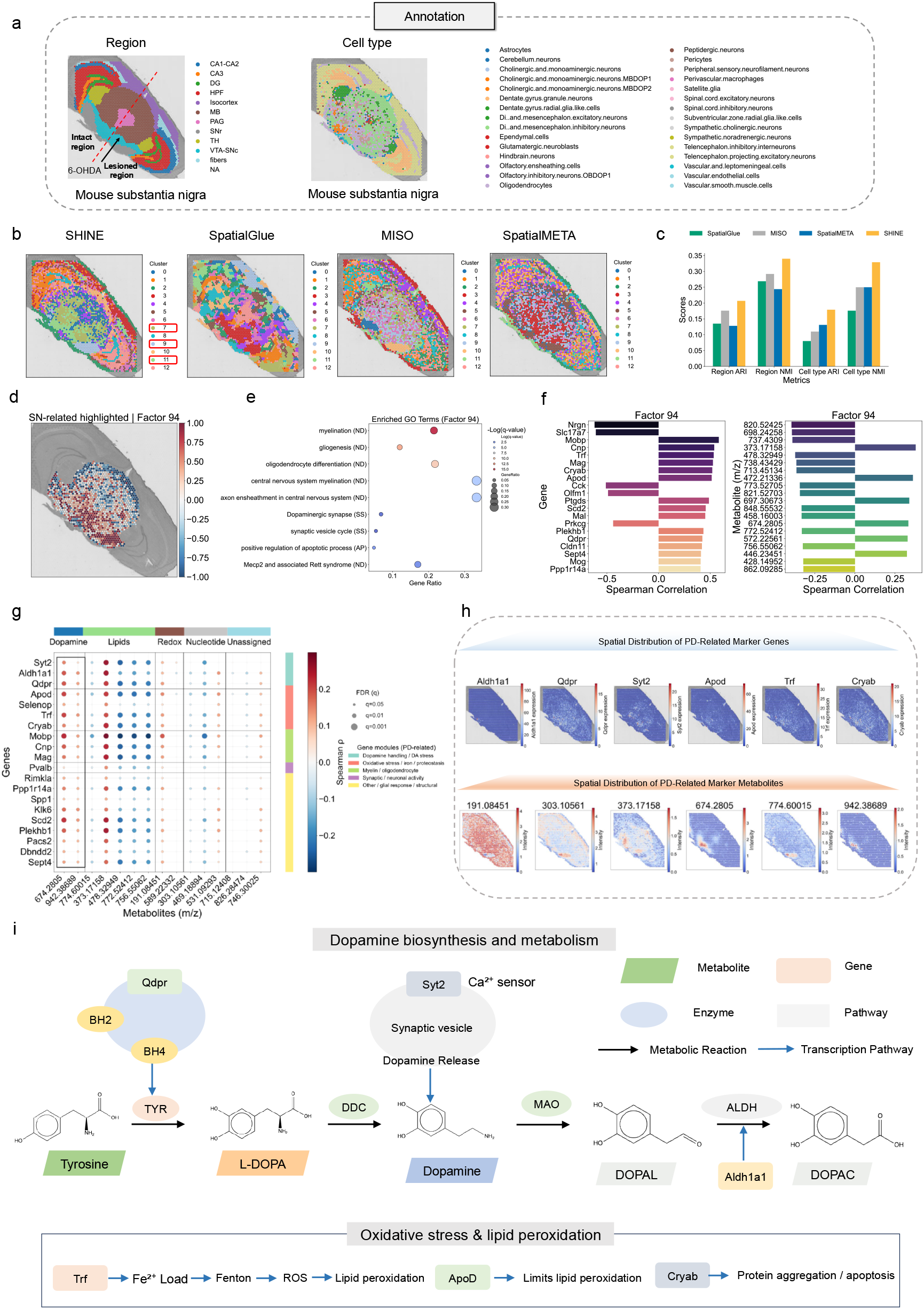
SHINE analysis of spatial metabolic–transcriptional microenvironments in the mouse SN. **a**, Region annotations and cell-type labels of the lesioned and intact hemispheres from unilateral 6-OHDA–treated mouse SN. **b**, Spatial clustering results from SHINE, MISO, SpatialGlue and SpatialMETA. SHINE delineates anatomically coherent domains, including dopaminergic neuron–depleted SNc regions, while baseline methods produce fragmented or misaligned boundaries. **c**, Clustering performance comparison using ARI and NMI across region annotations, cell types, and PD-related categories. **d**, Spatial distribution of latent Factor 94, enriched in the lesioned SNc. **e**, GO enrichment of Factor 94-associated genes, highlighting PD-relevant pathways such as dopamine metabolism, oxidative stress and neuroinflammation. **f**, Top genes and metabolites correlated with Factor 94. **g**, Gene–metabolite correlation map revealing coordinated metabolic–transcriptional modules associated with PD-related spatial domains. **h**, Spatial distribution of representative PD marker genes and metabolites within lesioned SNc microenvironments. **i**, Pathway-level reconstruction, integrating spatially mapped gene and metabolite features.

SHINE’s RNA-metabolite integration enabled the identification of several PD-associated clusters in the SN. Differential analysis between lesioned and control hemispheres highlighted clusters (e.g., Clusters 7, 9 and 11) with distinct PD-driven molecular perturbations (Supplementary Fig. 7).

SHINE also recovered coherent gene–metabolite correlation structures in the SN data, which allowed us to perform latent factor decomposition shared between genes and metabolites. As a prominent example, we identified Factor 94 as a PD-enriched latent axis with a striking spatial localization to the lesioned SNc (Supplementary Fig. 8). Notably, Factor 94 featured genes associated with oxidative stress (*Cryab, Sod2*), myelin and lipid metabolism (*Mobp, Mag*), and dopaminergic neuron vulnerability (*Aldh1a1, Qdpr*). These genes reflect core pathologies of PD, including glial stress responses, mitochondrial dysfunction, myelin disruption, and impaired BH4-mediated dopamine synthesis. This analysis identified Factor 94 as a PD-enriched latent axis with strong spatial localization to the lesioned SNc (Fig. 3d). Consistent with these molecular signatures, GO enrichment analysis of Factor 94–associated genes highlighted biological processes relevant to Parkinson’s disease, including dopamine metabolism, myelination-related pathways, oxidative stress responses and neuroinflammatory processes (Fig. 3e). The top-ranked genes and metabolites, identified by Spearman correlation with this factor, included multiple components implicated in dopaminergic function, oxidative stress and glial regulation (Fig. 3f).

Based on these results, a subset of marker genes and metabolites showing pronounced hemispheric asymmetry was selected from PD-associated clusters and from Factor 94–associated features and summarized in Supplementary Table S2. Gene–metabolite correlation patterns (Fig. 3g) revealed distinct coordinated molecular modules, reflecting parallel engagement of neuronal and glial processes. The spatial distributions of all genes and metabolites included in the hierarchical clustering analysis in Fig. 3g are shown in Supplementary Fig. 9. Notably, spatial contrasts were more pronounced for metabolite signals than for gene expression, which appeared comparatively subtler across tissue space. Representative PD-relevant genes and metabolites were further selected for focused visualization in Fig. 3h.

Integrating these signatures, SHINE delineated a set of coordinated molecular programs that together define the pathological landscape of Parkinson’s disease in the SN (Fig. 3i). Central to these programs was a dopaminergic–BH4 axis governing dopamine biosynthesis through regulation of tyrosine hydroxylase cofactor availability, thereby linking impaired dopamine production to selective neuronal vulnerability [20]. This core program was accompanied by a myelin–lipid program reflecting oligodendrocyte-associated membrane and myelin alterations, as well as an inflammation–energy program capturing astrocytic and microglial activation coupled with metabolic remodeling. In parallel, SHINE resolved an iron–oxidative program characterized by increased iron burden, reactive oxygen species generation and lipid peroxidation, consistent with ferroptosis-related stress in vulnerable dopaminergic neurons [21]. These molecular alterations converged with a PD-canonical program encompassing hallmark features of the disease, including markers of dopaminergic neuron loss, *α*-synuclein pathology and impaired protein homeostasis. Together, these interlinked gene–metabolite programs illustrate the multifactorial nature of PD pathogenesis, in which disrupted dopamine synthesis and handling are accompanied by metabolic alterations, including lipid and oxidative stress–related features, alongside glial responses and proteostasis imbalance, collectively shaping lesion-specific molecular microenvironments in the SN.

In conclusion, these results highlight SHINE’s ability to resolve spatial molecular microenvironments in neurodegenerative disease. We next extended the analysis to human spatial multi-omics datasets from lung and breast cancers to evaluate its generalizability across species, tissues and disease contexts.

### 2.4 SHINE’s integrative RNA-metabolite embeddings resolve spatial tumor microenvironments in human lung cancer

To examine the generalizability of SHINE across species and tissue contexts, we applied the framework to a human lung cancer spatial multi-omics dataset [10] (Fig. 4a). Using pathologist-defined annotations as reference, SHINE recovered most major tissue regions and additionally identified vascular structures, including blood vessels, that were less clearly resolved by baseline methods (Fig. 4b). Spatial clustering and UMAP projections further showed that SHINE more clearly delineated tumor boundaries and blood vessel structures, with substantially reduced fragmentation compared to baseline approaches. Quantitative evaluation confirmed higher agreement with pathologist annotations, as reflected by improved ARI and NMI scores across annotated cell types (Fig. 4c).

To assess whether the inferred spatial domains corresponded to biologically meaningful cellular niches, we quantified the cell-type composition of each cluster (Fig. 4d). Tumor-enriched clusters were dominated by malignant epithelial cells, indicating that SHINE resolves fine-grained microenvironmental boundaries within lung cancer tissue. Next, we investigated whether the molecular programs within these spatial clusters reflected canonical features of lung cancer biology. The tumor-associated clusters identified by SHINE exhibited elevated expression of epithelial and oncogenic marker genes typically upregulated in lung adenocarcinoma. In parallel, these same clusters showed enrichment of cancer-associated metabolic features such as various phosphatidylcholines and lipid-related ions within the malignant regions (Fig. 4e and Supplementary Table S3). This concordance between transcriptional and metabolic alterations indicates that each spatial cluster corresponds to a functionally specialized tumor or microenvironment niche within the lung adenocarcinoma tissue. Integrative analysis of spatial transcriptomic markers and metabolite features further revealed that SHINE’s clusters capture distinct aspects of tumor biology that extend beyond what either modality alone would show.

**Fig. 4.**
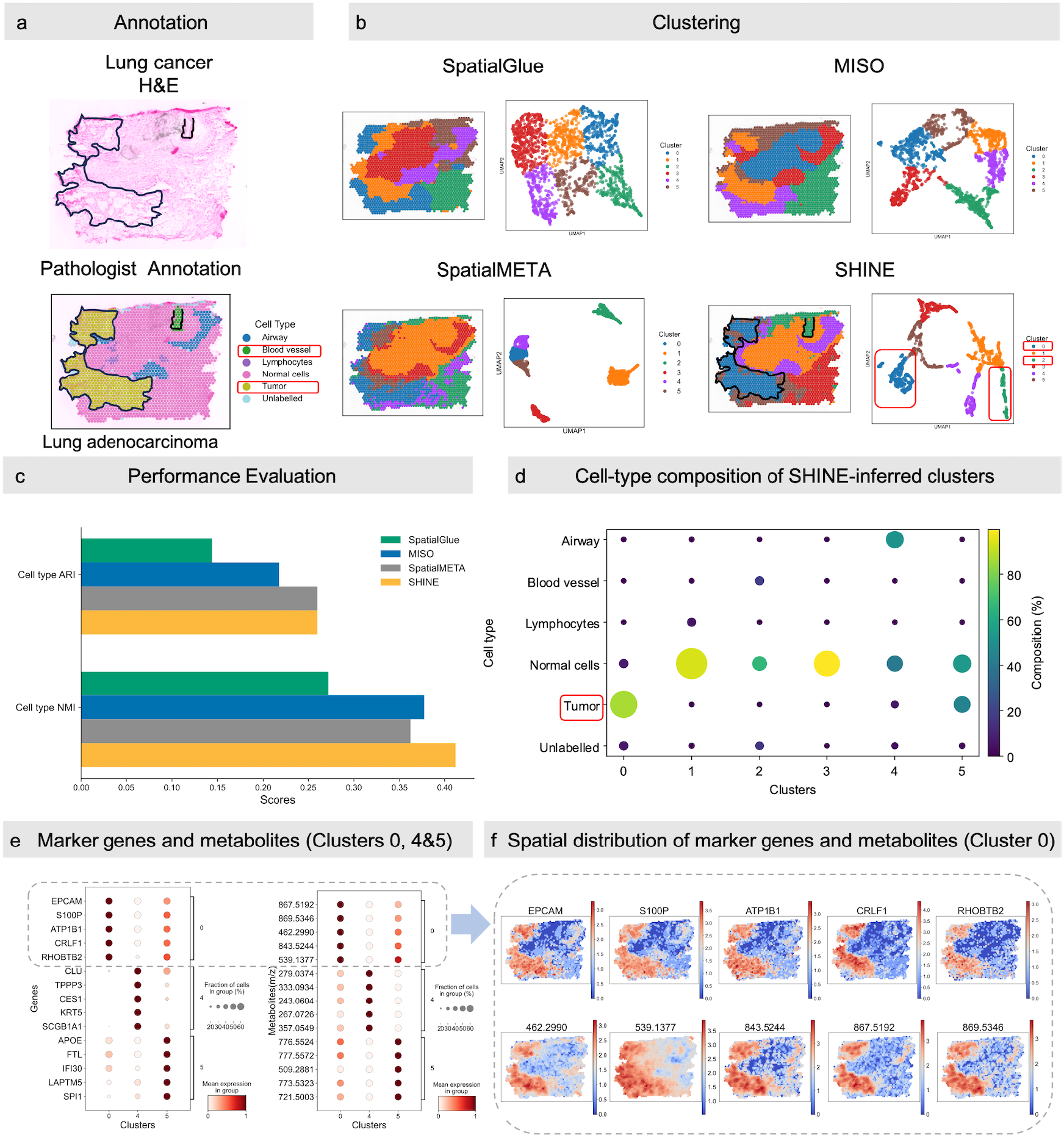
SHINE identifies spatial tumor–immune microenvironments in human lung cancer. **a**, H&E staining and pathologist-provided annotations of human lung cancer tissue. **b**, Spatial clustering results and UMAP projections generated by SpatialGlue, MISO, SpatialMETA and SHINE. **c**, Clustering accuracy comparison using ARI and NMI metrics based on pathologist annotations. **d**, Cell-type composition of SHINE-inferred clusters. **e**, Marker genes and metabolites across tumor-core and neighboring clusters (Clusters 0, 4 and 5). **f**, Spatial distribution of marker genes and metabolites across tumor-core cluster (Cluster 0).

Cluster 0 represented the tumor core, marked by elevated expression of canonical epithelial tumor markers (e.g., *EPCAM*and *S100P*). Consistent with this annotation, spatial visualization revealed that these genes were strongly enriched within the tumor core region (Fig. 4f). Several metabolic features observed in this region, including highly unsaturated phosphatidylglycerols such as PG(42:9) (*m/z* 843.5244) and PG(42:11) (*m/z* 867.5192), are consistent with previously reported spatial metabolite–transcript associations in lung cancer tissue [10]. Notably, the glycogen-associated metabolite maltotriose (*m/z* 539.1377) displayed a spatial distribution extending beyond the epithelial tumor core, indicating metabolically influenced transition zones surrounding the tumor and suggesting coordinated tumor–stroma metabolic interactions.

Surrounding the tumor core of Cluster 5, SHINE resolved a peritumoral immune–stromal interface characterized by lipid remodeling signatures, including ether-linked phosphatidylethanolamines such as PE(P-40:5) (*m/z* 776.5524) and lysophospholipids exemplified by LysoPG(18:1) (*m/z* 509.2881). These metabolic features spatially aligned with immune- and lysosome-associated gene expression (e.g., *FTL, IFI30, SPI1*), indicating an invasion-associated microenvironment involving lipid handling, iron metabolism, and antigen processing rather than intrinsic tumor-cell biosynthesis.

In contrast, an adjacent airway-associated epithelial compartment (Cluster 4) marked by *SCGB1A1* and *KRT5* exhibited enrichment of nucleoside metabolism, exemplified by uridine (*m/z* 279.0374) and inosine (*m/z* 267.0726), together with lysophospholipid signaling such as LysoPA(8:0) (*m/z* 333.0934). While nucleoside-associated metabolic signatures have been previously noted in lung spatial multi-omics datasets, SHINE clearly segregates this preserved epithelial niche from tumor-associated metabolic remodeling, highlighting structured transcriptional–metabolic organization across tumor, stromal, and airway epithelial regions within the same tissue section.

In summary, these results demonstrate that SHINE resolves spatially segregated malignant, stromal and airway epithelial niches, each defined by distinct yet coordinated transcriptional and metabolic programs. Together, these results suggest that lung adenocarcinoma transcriptional programs are coupled to coordinated metabolic reprogramming, linking proliferative epithelial states with spatially extended lipid and glycogen metabolism that shapes the surrounding tumor microenvironment.

### 2.5 Spatial characterization of tumor microenvironment in human breast cancer tissues using SHINE

To further evaluate the robustness of SHINE in studying cancers, we applied the framework to spatial paired SM and ST data from human breast cancer tissue sections [10]. Tumor and stromal compartments defined by pathologist annotations are shown together with hematoxylin and eosin (H&E) staining and pathology-guided single-cell-derived labels, which serve as ground truth references (Fig. 5a; see more in Methods). Compared with SpatialGlue, MISO and SpatialMETA, SHINE produced sharper tumor–stroma boundaries and more faithfully resolved localized stromal heterogeneity and intratumoral subtype variation (Fig. 5b), resulting in consistently higher agreement with both pathologist annotations and single-cell–derived labels (Fig. 5c).

**Fig. 5.**
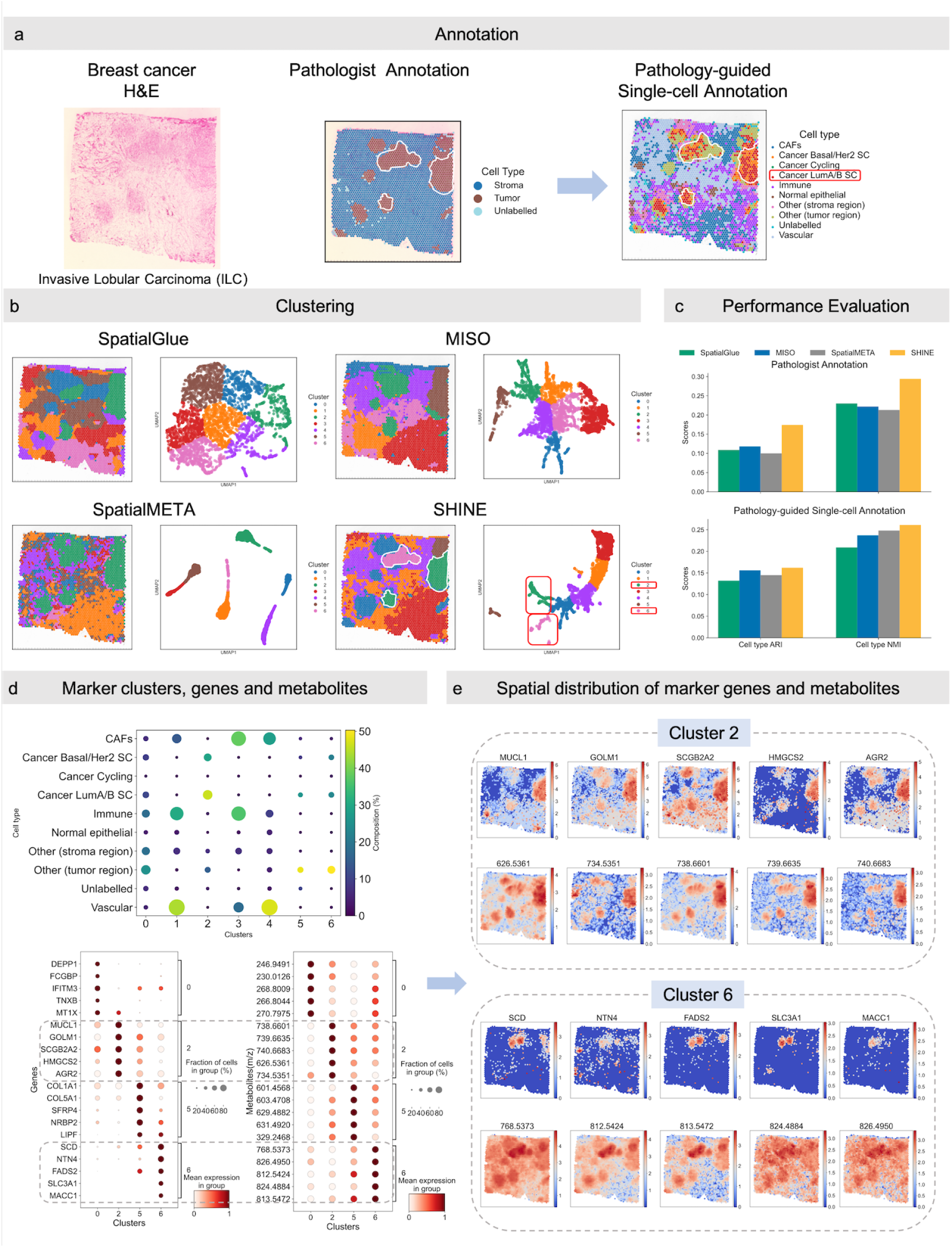
SHINE delineates tumor–stroma architecture and subtype-specific programs in human breast cancer. **a**, H&E staining and pathologist annotations of human breast cancer tissue, highlighting tumor and stromal regions (left, center). Right: pathology-guided single-cell annotation reveals cellular diversity across tumor and stromal compartments. **b**, Spatial clustering and UMAP projections from SHINE, MISO, SpatialGlue and SpatialMETA. SHINE captures sharp tumor–stroma boundaries and stromal heterogeneity, with more distinct subtype separation than baseline models. **c**, Clustering performance comparison using ARI and NMI with two benchmarks: pathologist annotations (left) and single-cell–derived reference labels (right). **d**, Cluster-specific marker genes and metabolites defining tumor-associated domains. **e**, Spatial distribution of tumor-associated clusters(Clusters 2 and 6) marker genes and metabolites.

To determine whether the inferred spatial domains corresponded to biologically meaningful tumor states, we examined their transcriptional and metabolic programs. Tumor-enriched regions exhibited high expression of luminal epithelial markers, consistent with hormone receptor–positive breast cancer (Fig. 5d). These regions were further characterized by elevated levels of lipid metabolites implicated in tumor progression and membrane remodeling (Fig. 5e) (Supplementary Table S4), which have been implicated in lipid storage, membrane biogenesis and hormone-responsive growth in luminal breast tumors [22–24].

At higher resolution, SHINE resolved multiple spatially organized tumor and microenvironmental niches by Clusters 0, 2, 5, and 6 (Fig. 5d), each with a set of uniquely enriched genes and metabolites (Fig. 5d). Among the metabolites discussed in this analysis, PS(38:3) (*m/z* 812.5424) was previously annotated in the original study of this dataset [10]. The remaining metabolites were putatively assigned through accurate mass matching to entries in the Human Metabolome Database (HMDB) and were not explicitly emphasized in the prior correlation-based analysis. Thus, the novelty of SHINE does not lie in the identification of new molecular species, but in organizing annotated metabolites and transcripts into spatially coherent, higher-order programs.

Specifically, Cluster 2 showed strong concordance with Luminal A/B–like regions identified by single-cell projection and displayed coordinated upregulation of luminal gene signatures (*MUCL1, GOLM1, SCGB2A2, HMGCS2* and *AGR2*) together with enrichment of cholesterol esters (CE(20:0), *m/z* 738.6601/739.6635/740.6683) [25], ether-linked phospholipids (PC(O-30:1), *m/z* 734.5351) [26], and ceramides (Cer(d38:2), *m/z* 626.5361) [27], defining a secretory, hormone-responsive tumor core. Adjacent to this core, Cluster 6 marked a metabolically distinct tumor substate characterized by elevated expression of lipid desaturation enzymes (*SCD, FADS2*) and broad enrichment of membrane phospholipids and oxidized phosphatidylethanolamines, consistent with enhanced membrane biogenesis and oxidative lipid remodeling [28]. The partially segregated yet spatially adjacent distributions of Clusters 2 and 6 indicate spatially organized intra-tumoral heterogeneity, with a luminal secretory tumor core transitioning toward lipid-remodeled tumor regions. Representative spatial distributions of marker genes and metabolites for Clusters 2 and 6 are shown in Fig. 5e.

In contrast, Cluster 5 localized to stromal regions enriched for extracellular matrix and fibroblast-associated genes (*COL1A1, COL5A1, SFRP4, NRP2*) and showed prominent accumulation of diacylglycerols and polyunsaturated fatty acid–related metabolites, indicative of an actively remodeling CAF-rich stromal niche [29]. Surrounding these domains, Cluster 0 formed a peritumoral boundary enriched for interferon-responsive and stress-associated genes (*IFITM3, MT1X, DEPP1, TNXB* and *FCGBP*) together with sulfated catecholamine and phenolic metabolites, consistent with an exposure- and inflammation-associated tumor interface. Spatial distribution of marker genes and metabolites for Clusters 0 and 5 is provided in Supplementary Fig. 10.

In summary, these results demonstrate that high-order spatial multi-omic integration reveals coordinated transcriptional and lipid metabolic programs across tumor and microenvironmental compartments. Beyond recapitulating major tumor–stroma organization, SHINE resolves subtype-associated molecular programs within the tumor, highlighting the value of high-order spatial modeling for uncovering functional heterogeneity in breast cancer tissues.

## 3 Discussion

In this study, we present SHINE, a hypergraph-based framework for integrative analysis of spatial metabolomics and transcriptomics that explicitly models higher-order cross-modality interactions within tissue microenvironments. By moving beyond pairwise associations and implicit latent representations, SHINE captures coordinated metabolic–transcriptional programs that emerge from the collective organization of transcripts, metabolites and spatial context, providing a principled framework for modeling higher-order organization in spatial multi-omics data. Across diverse datasets spanning species, tissues and disease settings, SHINE consistently recovered biologically coherent spatial domains and outperformed existing integration strategies, demonstrating its robustness and general applicability for spatial multi-omics analysis.

The ability to model higher-order spatial interactions has important implications for understanding complex tissue phenotypes. Increasing evidence suggests that spatially structured diseases, including cancer and neurodegeneration, are shaped not by isolated molecular events but by coordinated programs linking transcriptional regulation, metabolic state and cellular composition across tissue microenvironments. From a spatial metabolomics perspective, particularly when combined with transcriptomics, this integrative view enables the identification of metabolic niche organization within tumors, including tumor–stroma interfaces, immune-associated metabolic gradients and subtype-specific lipid remodeling programs that are difficult to resolve using transcriptomics alone.

The performance gains of SHINE likely arise from its explicit modeling of higher-order multi-modal interactions through hypergraph-based representation learning. By jointly integrating spatial transcriptomics, metabolite abundance profiles and spatial neighborhood structure, the framework captures coordinated molecular programs distributed across multiple spatial scales rather than relying on pairwise correlations or single-modality clustering. The input to SHINE consists of spatially aligned multi-omics features and spatial coordinates, whereas the output includes integrated embeddings, spatial domain segmentation and interpretable gene–metabolite programs, enabling systematic characterization of tissue microenvironment organization. This modeling strategy is broadly applicable across biological contexts. In tumor tissues, it facilitates the delineation of metabolically heterogeneous tumor niches and immune–stromal interfaces; in neurodegenerative models, it supports the identification of spatially localized metabolic–transcriptional alterations associated with neuronal vulnerability and inflammation. Together, these findings highlight the value of higher-order spatial multi-omics modeling for revealing systems-level molecular organization across diverse disease settings.

Despite these advances, several limitations should be considered. SHINE is designed to integrate spatially resolved measurements that capture comparable tissue contexts across modalities, and its performance may depend on the quality of the underlying spatial correspondence, particularly in multi-section integration settings. In addition, while the hypergraph formulation provides a flexible representation of higher-order structure, defining biologically optimal hyperedge construction strategies remains an open challenge and may benefit from incorporating prior biological knowledge, adaptive learning schemes, or data-driven discovery of higher-order structure. Future extensions of SHINE could integrate additional modalities, such as spatial proteomics or epigenomics, and more explicitly bridge single-cell and tissue-level data to support multiscale analysis. Together, these developments may further enhance the utility of SHINE as a general platform for decoding complex tissue organization and disease-associated microenvironments.

## 4 Methods

### 4.1 Spatial Metabolomics and Transcriptomics Data Alignment

#### 4.1.1 Datasets Collection

SHINE was evaluated using paired spatial metabolomics and spatial transcriptomics datasets derived from both mouse and human tissues (Fig. 1A). Spatial omics data from the mouse PD model were obtained from Vicari et al. [8]. In this model, coronal brain sections encompassing the striatum and SN were collected from mice subjected to unilateral 6-OHDA lesions, together with matched control animals, resulting in selective dopaminergic neuron degeneration in the lesioned hemisphere. Human lung and breast cancer spatial omics datasets were derived from Godfrey et al. [10] and consisted of surgical resections containing both malignant regions and adjacent non-malignant tissues. Due to the relatively limited cellular diversity in the breast cancer samples, we further refined cell-type annotations at the single-cell level, guided by pathologist-provided annotations, with single-cell annotation data obtained from [30]. Spatial metabolomics and spatial transcriptomics profiling was performed on adjacent or serial tissue sections to enable spatially matched multi-omics analysis.

Spatial metabolomics data were acquired using MSI platforms adapted to species- and tissue-specific experimental designs. For the mouse PD model, spatial metabolomics profiling was performed using MALDI–MSI, employing either FMP-10 or DHB as matrices targeting different metabolite classes. MALDI–MSI generated high-resolution ion intensity maps across a regular spatial pixel grid for each detected *m/z* feature. Spatial transcriptomics profiling was subsequently performed on the same tissue sections using the 10x Genomics Visium platform, which captures spot-level gene expression profiles (55 µm spot diameter; 100 µm center-to-center spacing). RNA detectability following MALDI–MSI acquisition has been previously validated, enabling spatially matched multi-omics measurements.

For human lung and breast cancer samples, spatial metabolomics data were generated using desorption electrospray ionization mass spectrometry imaging (DESI–MSI) at an approximate spatial resolution of 100 µm per pixel without matrix application. Corresponding spatial transcriptomics data were obtained using the 10x Genomics Visium platform, and sequencing libraries were processed using the standard Space Ranger pipeline to generate normalized spot-by-gene expression matrices. The comparable spatial resolutions of MSI and Visium measurements enabled downstream spatial integration across modalities.

#### 4.1.2 Data Alignment

For the mouse PD dataset, spatial metabolomics and spatial transcriptomics measurements were acquired sequentially on the same tissue section but in different experimental modalities. As a result, spatial co-registration was required to align MSI pixels with Visium spots. High-resolution H&E images were used as anatomical references (Fig. 1A). Spatial alignment was performed by first applying rigid transformations, including translation, rotation and scaling, to achieve global alignment between MSI data and Visium coordinates, followed by TPS transformations to account for local nonlinear tissue deformations. This procedure enabled spatial correspondence between MSI pixels and Visium spots.

For the human lung and breast cancer datasets, spatial metabolomics and spatial transcriptomics data were also obtained from the same tissue section and were provided with pre-aligned spatial coordinates. Therefore, no additional alignment procedures were applied, and the provided alignments were used directly for all downstream analyses. When an exact one-to-one correspondence could not be achieved during alignment, each Visium spot was associated with its nearest MSI pixel in physical space. Metabolites were matched based on shared molecular identities inferred from common *m/z* features. MSI intensities were normalized (for example, using total-ion current normalization) and log-transformed, while Visium gene expression counts were normalized and log-transformed following the standard spatial transcriptomics workflow. These paired representations were subsequently used as input for Spatial Hypergraph Integration of Metabolic Networks and Gene Expression.

### 4.2 Spatial Hypergraph Integration of Metabolic Networks and Gene Expression

SHINE is a hierarchical spatial omics integration framework designed to jointly model SM and ST data across aligned tissue locations (Fig. 1B). The framework consists of four main stages: Spatial Omics Data Input, Intra-Modality Integration, Cross-Modality Hypergraph Integration and Decoding & Reconstruction.

#### 4.2.1 Spatial Omics Data Input

For each tissue section, SHINE operates on spatially aligned multi-omics measurements defined over a common set of spatial locations. Let *N* denote the number of spatial locations after multi-omics co-registration, *D*_*m*_ denote the dimensionality of metabolomic features, and *D*_*t*_ denote the dimensionality of transcriptomic features. For each spatial location *i*, SHINE takes as input a metabolomic feature vector 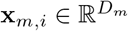, a transcriptomic feature vector 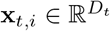, and a corresponding spatial coordinate vector 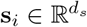.

The spatial coordinates **s**_*i*_ represent aligned positions obtained after spatial co-registration between spatial metabolomics and spatial transcriptomics. These aligned coordinates are used to construct a spatial neighborhood graph, in which nodes correspond to spatial locations and edges connect spatially proximal locations based on KNN. The resulting spatial graph is shared across modalities and provides the structural backbone for subsequent graph convolution and graph transformer operations.

Metabolomic and transcriptomic feature matrices are processed independently in the initial encoding stage but remain linked through their shared spatial coordinates. This formulation enables SHINE to preserve modality-specific feature spaces while ensuring that information integration occurs across spatially matched locations.

#### 4.2.2 Intra-Modality Integration

Within each omics modality, SHINE learns spatial representations by jointly modeling local spatial topology and feature-level molecular similarity across aligned tissue locations. Two complementary graphs are constructed over the same set of spatial nodes, including a spatial neighborhood graph derived from physical coordinates and a feature similarity graph capturing molecular relationships between locations. On the spatial graph *G*, local tissue structure is modeled through a SpatialGCN message passing scheme. For each spatial location *i*, modality-specific molecular measurements (that is, **x**_*t,i*_ for transcriptomics or **x**_*m,i*_ for metabolomics) and spatial coordinates **s**_*i*_ are first independently projected into a shared latent space and combined to form the initial node representation:

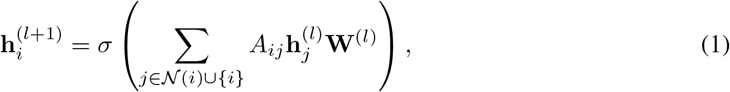

where *A*_*ij*_ denotes entries of the spatial adjacency matrix, *ϕ*(·) denotes a nonlinear activation function, and **W**_*f*_ and **W**_*s*_ are learnable projection matrices for molecular features and spatial coordinates, respectively. The resulting representation initializes the node embedding 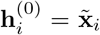, after which spatial propagation updates node representations according to:

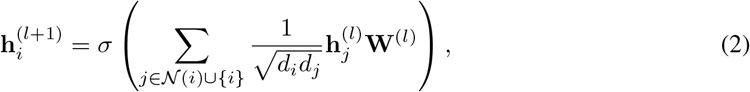

where *N* (*i*) denotes spatial neighbors of node *i* and *d*_*i*_ = |*N* (*i*) ∪ { *i* }| represents node degree after adding self-loops. This operation aggregates information from spatially adjacent locations, enabling embeddings to capture locally coherent tissue organization.

In parallel, long-range molecular dependencies are modeled using a Graph Transformer encoder operating on the feature similarity graph. Let **h**_*i*_ denote the input embedding for node *i*. For each attention head *h* = 1, …, *H*, pairwise attention scores between nodes *i* and *j* are computed as:

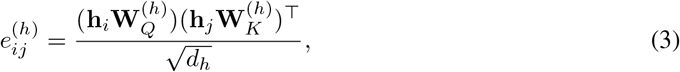

where 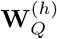 and 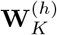 denote learned projection matrices and *d*_*h*_ represents the dimensionality of each attention head. Attention is constrained by graph connectivity by masking attention scores using the feature adjacency matrix, such that only connected nodes contribute to aggregation. Normalized attention weights are then obtained as

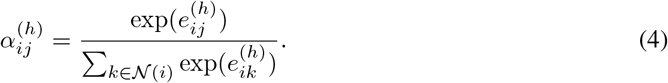

Each attention head produces an updated representation:

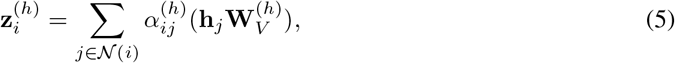

and multi-head outputs are concatenated and linearly transformed to obtain

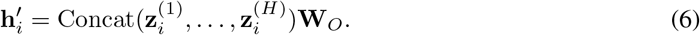

The spatial representation obtained from neighborhood propagation and the feature-aware representation learned through Graph Transformer attention are subsequently integrated through an attention-based fusion mechanism that adaptively balances local spatial structure and global molecular similarity. The resulting modality-specific embeddings are denoted as:

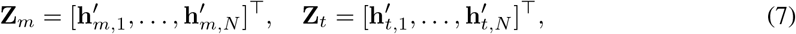

representing metabolomic and transcriptomic spatial embeddings, respectively.

These modality-specific embeddings encode spatial organization within each omics layer and serve as inputs for subsequent cross-modality integration.

#### 4.2.3 Cross-Modality Hypergraph Integration

To perform cross-modality integration, SHINE employs a directed causal attention mechanism, in which transcriptomic embeddings guide the aggregation of metabolomic representations. Given modality-specific spatial embeddings **Z**_*t*_ and **Z**_*m*_ obtained from the preceding intra-modality integration stage, directed cross-modality fusion produces a causal embedding:

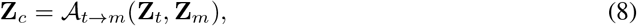

where *A*_*t → m*_(·) denotes a directed cross-modality attention operation mapping transcriptomic representations to metabolomic features.

In this formulation, each spatial location (spot) is treated as a node in the hypergraph. Let *V*_*H*_ = {*v*_1_, …, *v*_*N*_ } denote the set of spatial nodes shared across modalities, where *N* is the number of aligned spatial locations. Each node *v*_*i*_ is associated with a multi-modal embedding 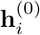 initialized from the causal cross-modality embedding matrix **Z**_*c*_, such that **H**^(0)^ = **Z**_*c*_. Hyperedges are constructed using a kNN strategy in the learned multi-modal embedding space. Specifically, each spatial node defines a hyperedge connecting itself and its neighboring spots:

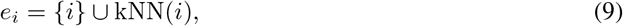

which represents a local spatial microenvironment composed of molecularly or spatially similar locations. Let 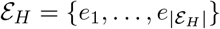 denote the hyperedge set. The resulting hypergraph structure is encoded by an incidence matrix:

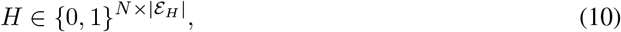

where *H*_*i,e*_ = 1 indicates that spatial node *i* belongs to hyperedge *e*. Using **Z**_*c*_ as initialization, hypergraph message passing is performed through node–hyperedge–node propagation to enable higher-order aggregation. At layer *l*, node embeddings are updated as:

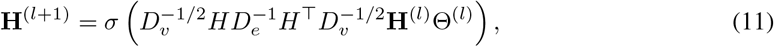

where 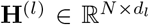 denotes node embeddings at layer *l, D*_*v*_ and *D*_*e*_ are diagonal matrices of node degrees and hyperedge sizes, respectively, and Θ^(*l*)^ is a learnable transformation matrix. After *L* hypergraph propagation layers, we obtain the final hypergraph-refined embedding matrix:

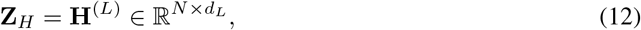

which represents the higher-order multi-modal spatial representation of all locations. This embedding serves as the unified latent representation used for downstream reconstruction and optimization.

This operation aggregates information across groups of spatial locations rather than pairwise neighbors, allowing spatial spots that share common microenvironmental context to exchange information through shared hyperedges. Stacking multiple hypergraph layers progressively refines multi-modal spot embeddings, enabling SHINE to capture higher-order spatial organization of joint transcriptomic–metabolomic programs.

#### 4.2.4 Decoding & Reconstruction

After cross-modality causal attention and hypergraph propagation, SHINE obtains a unified hypergraph-refined embedding matrix **Z**_*H*_ = [**z**_*H*,1_, …, **z**_*H,N*_]^*⊤*^, which captures higher-order spatial organization and directed transcriptomic-to-metabolomic interactions across tissue locations. This representation serves as the shared latent space for modality-specific decoding and reconstruction.

To recover modality-specific observations, SHINE first applies modality-specific spatial decoders that propagate latent embeddings over spatial graphs to generate modality-aware predictions:

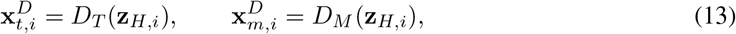

where *D*_*T*_ (·) and *D*_*M*_ (·) denote GCN-based decoders that enforce consistency with local tissue topology through spatial message passing.

In parallel, graph-transformer reconstruction modules further project latent embeddings back to the original feature spaces:

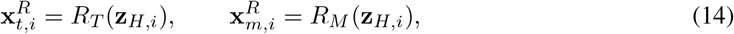

where *R*_*T*_ (·) and *R*_*M*_ (·) denote transformer-based reconstruction networks preserving modality-level semantic information learned during feature encoding. The corresponding reconstruction outputs are denoted as 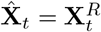 and 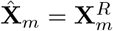, while decoding outputs are denoted as 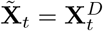 and 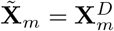.

The overall optimization objective jointly minimizes decoding and reconstruction errors together with cross-modality alignment induced by directed causal attention. The total loss is defined as:

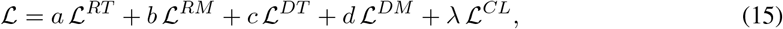

Where

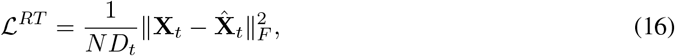

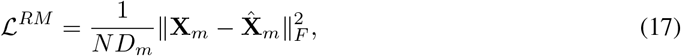

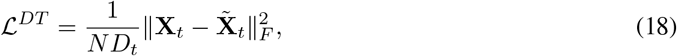

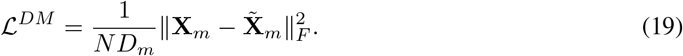

To further preserve transcriptomic–metabolomic correspondence established through directed causal attention, SHINE incorporates a contrastive alignment objective defined on modality-aware projections derived from the unified latent representation **Z**_*H*_. Specifically, modality-specific projections are obtained as:

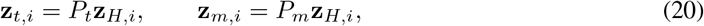

where *P*_*t*_ and *P*_*m*_ denote learnable projection functions for transcriptomic and metabolomic modalities. The contrastive objective is defined as:

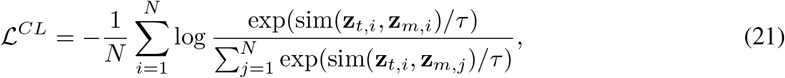

where sim(·,·) denotes cosine similarity and *τ* is a temperature parameter. Joint optimization of these objectives enables SHINE to learn spatially coherent, causally informed, and cross-modality aligned representations for downstream analysis.

## Acknowledgements

We thank the members of the StatBiomed Lab for their support throughout this study, and Dr. Lingyu Li and Jiamu James Qiao for their valuable contributions to discussions and code validation. This study was jointly supported by the National Natural Science Foundation of China (Grant No. 62502385), Hong Kong Scholars Program (Grant No. XJ2024050), the Research Grants Council of the Hong Kong SAR, China (Grant No. T12-705-24-R and 17126725).

## Declarations

### Code availability

Source code of this study is available at: https://github.com/Elsa-bingxue/SHINE

### Author contribution

Y.H. and B.D. conceived the study; B.D. implemented the SHINE framework and performed all benchmarking analyses. Y.H. and J.W. supervised the study and provided conceptual guidance; B.D. and Y.H. wrote the manuscript, with critical revisions from J.W. All authors approved the final manuscript.

