## Supplementary material for "SHINE: Decoding transcriptional-metabolic microenvironments through higher-order spatial integration": /: Supplementary.pdf

### Supplementary Methods

##### FMP-10 derivatization and mass spectrometry annotation

Metabolites in mouse striatum and substantia nigra brain slides were analyzed using FMP-10 derivatization–assisted mass spectrometry. The proton mass was set to $m_{\mathrm{H}} = 1.007276$ Da. The equivalent mass increment of a single FMP-10 derivatization was $\Delta = 268.11229$ Da. Common neutral losses, including $CH_3$ (15.02347 Da) and $H_2O$ (18.01056 Da), were considered during annotation.

Neutral monoisotopic masses ($M$) were back-calculated from observed $m/z$ values according to derivatization patterns, including single and double derivatization events with corresponding neutral losses. Theoretical adduct masses were calculated for $[M+H]^+$, $[M+Na]^+$, $[M+Li]^+$ and $[M+NH_4]^+$. Mass spectra were calibrated using $m/z$ 715.59 as the lock mass.

##### Spatial multi-omics integration and latent factor analysis

Spatial transcriptomic and metabolomic data were integrated using the SHINE framework. Latent representations were extracted for each modality and jointly optimized through hypergraph-based integration. Molecular features associated with latent factors were identified using Spearman correlation analysis, and spatial distributions were visualized for biological interpretation.

##### Implementation details

SHINE was implemented in PyTorch and run on CPU. The latent dimension was set to 256 and models were trained for 300 epochs using Adam (learning rate $1 \times 10^{-4}$, weight decay $1 \times 10^{-3}$). Feature graphs were constructed using $k$-nearest neighbors ($k = 20$), and hyperedges were constructed using $k$-nearest neighbors ($k = 10$) on concatenated transcriptomic and metabolomic features. Spatial graphs were constructed from aligned coordinates using a distance threshold defined as the 5th percentile of pairwise distances. The training objective combined reconstruction, correspondence and InfoNCE contrastive losses (temperature $T = 0.07$), with initial weights $(2, 2, 1, 1)$.

##### Spatial alignment and data preprocessing for benchmarking

For fair comparison across methods, identical spatial alignment and preprocessing procedures were applied to SHINE and benchmark models. Tissue sections were first aligned using rigid registration followed by non-rigid refinement using thin-plate spline transformation (Fig. 1A). Transcriptomic profiles were restricted to the top 2,000 highly variable genes, whereas metabolomic profiles retained all detected metabolites after quality control.

All benchmark methods were evaluated on identical aligned datasets and feature sets. Human lung and breast cancer datasets were provided in pre-aligned form and were directly used for evaluation. For SpatialMETA, preprocessing followed the original publication, and results are shown in Supplementary Fig. 11.

#### Supplementary Figures

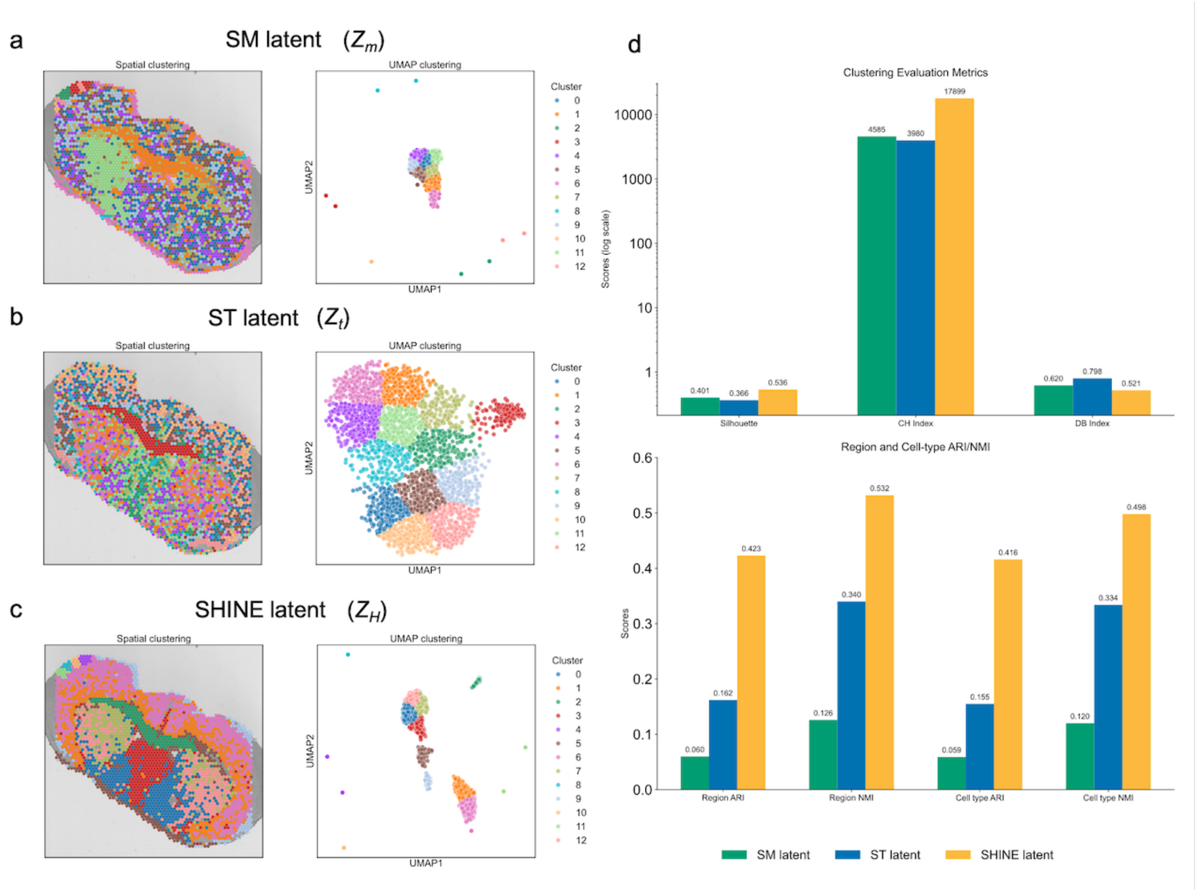


**Supplementary Fig. 1 | The importance of multi-omics integration. a–c**, Spatial and UMAP clustering using metabolomic ($Z_m$), transcriptomic ($Z_t$), and integrated SHINE ($Z_H$) embeddings, respectively. **d**, Quantitative evaluation of clustering performance and biological concordance using internal metrics together with region/cell-type ARI and NMI scores.

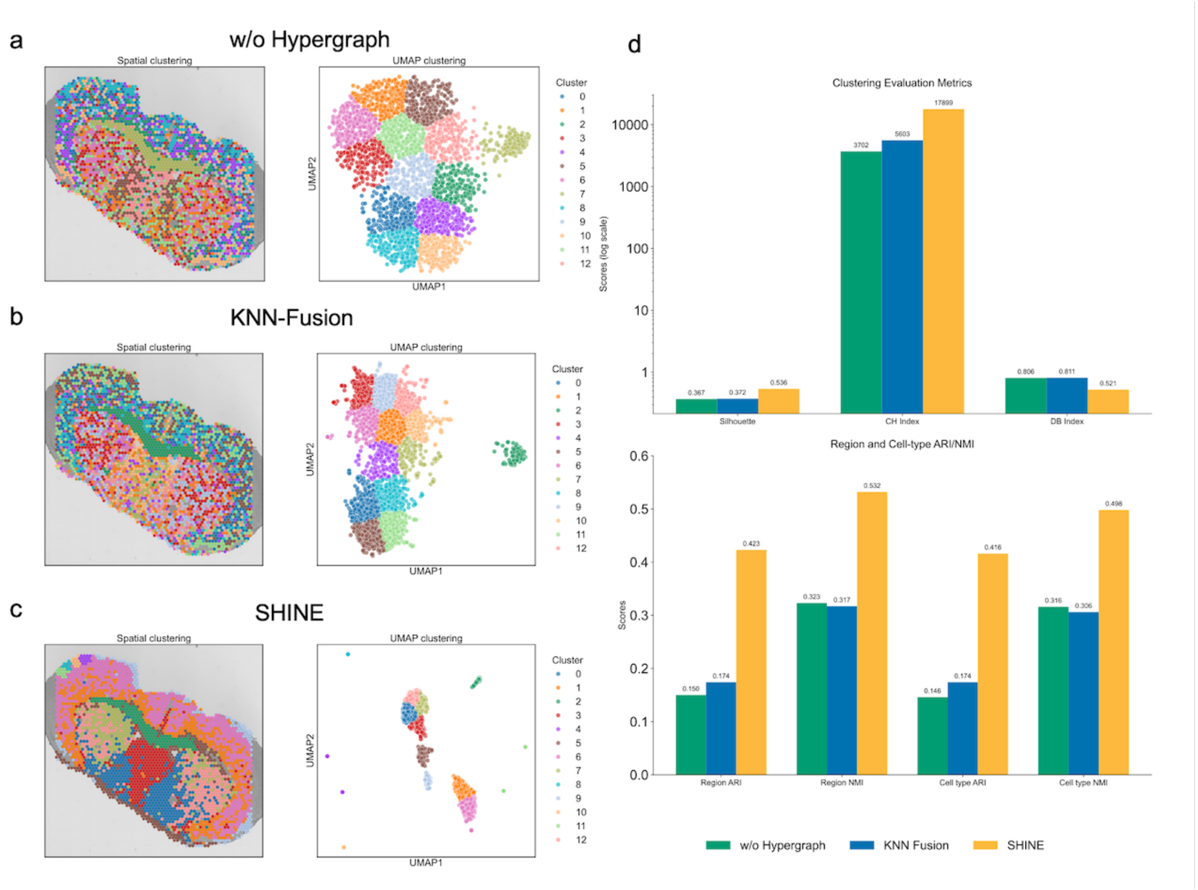


**Supplementary Fig. 2 | Ablation study to validate the contribution of each component to the performance of the SHINE. a–c**, Spatial and UMAP-based clustering results of model variants w/o Hypergraph (removing the hypergraph-based integration), KNN-Fusion (replacing hypergraph-based integration with KNN-based fusion), and the full SHINE; **d**, Quantitative comparison of the three settings using internal clustering metrics and region/cell-type ARI/NMI.

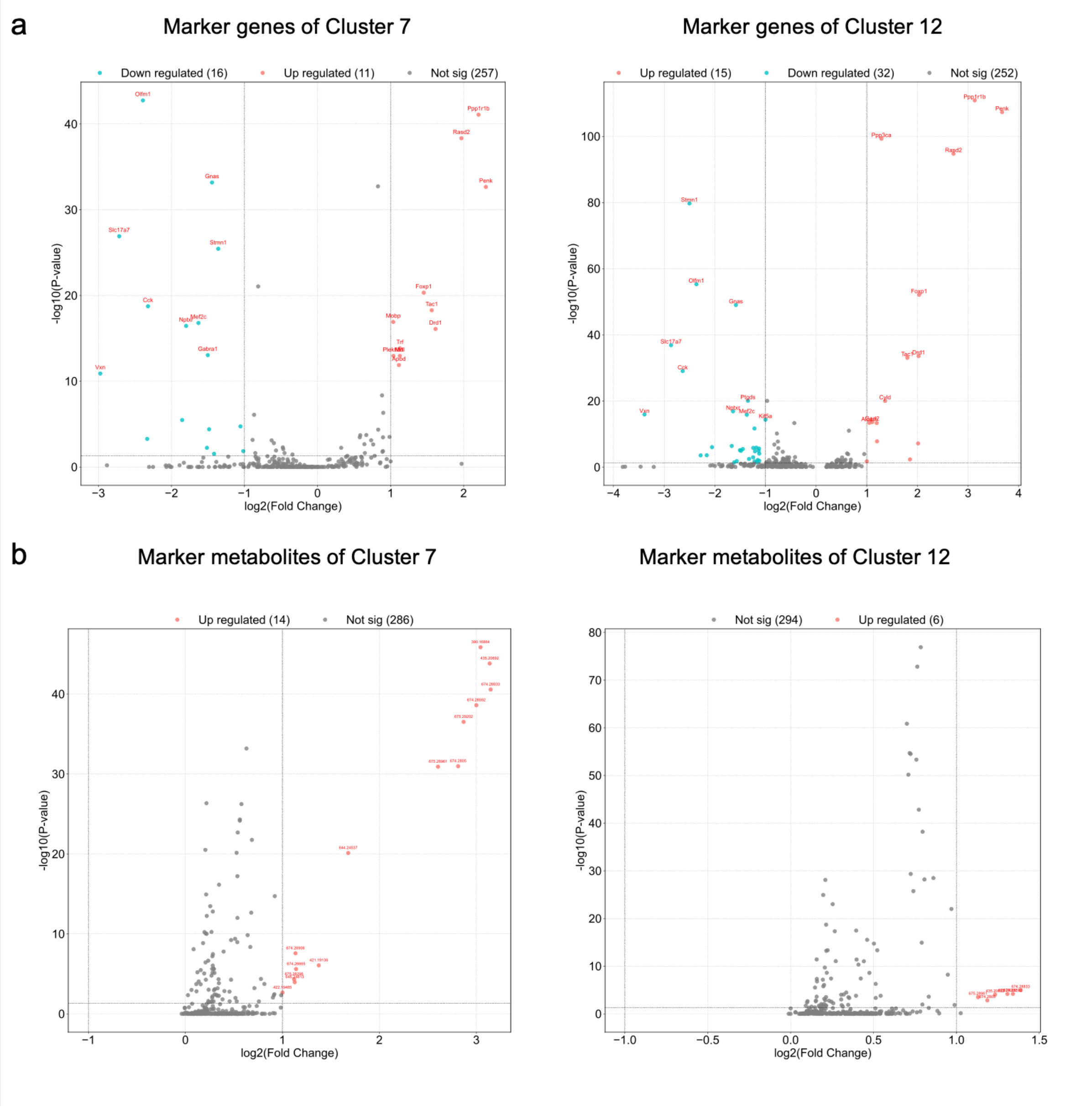


**Supplementary Fig. 3 | Marker genes and metabolites in Cluster 7 and Cluster 12. a**, Differentially expressed marker genes of representative clusters; **b**, Differentially abundant marker metabolites of representative clusters.

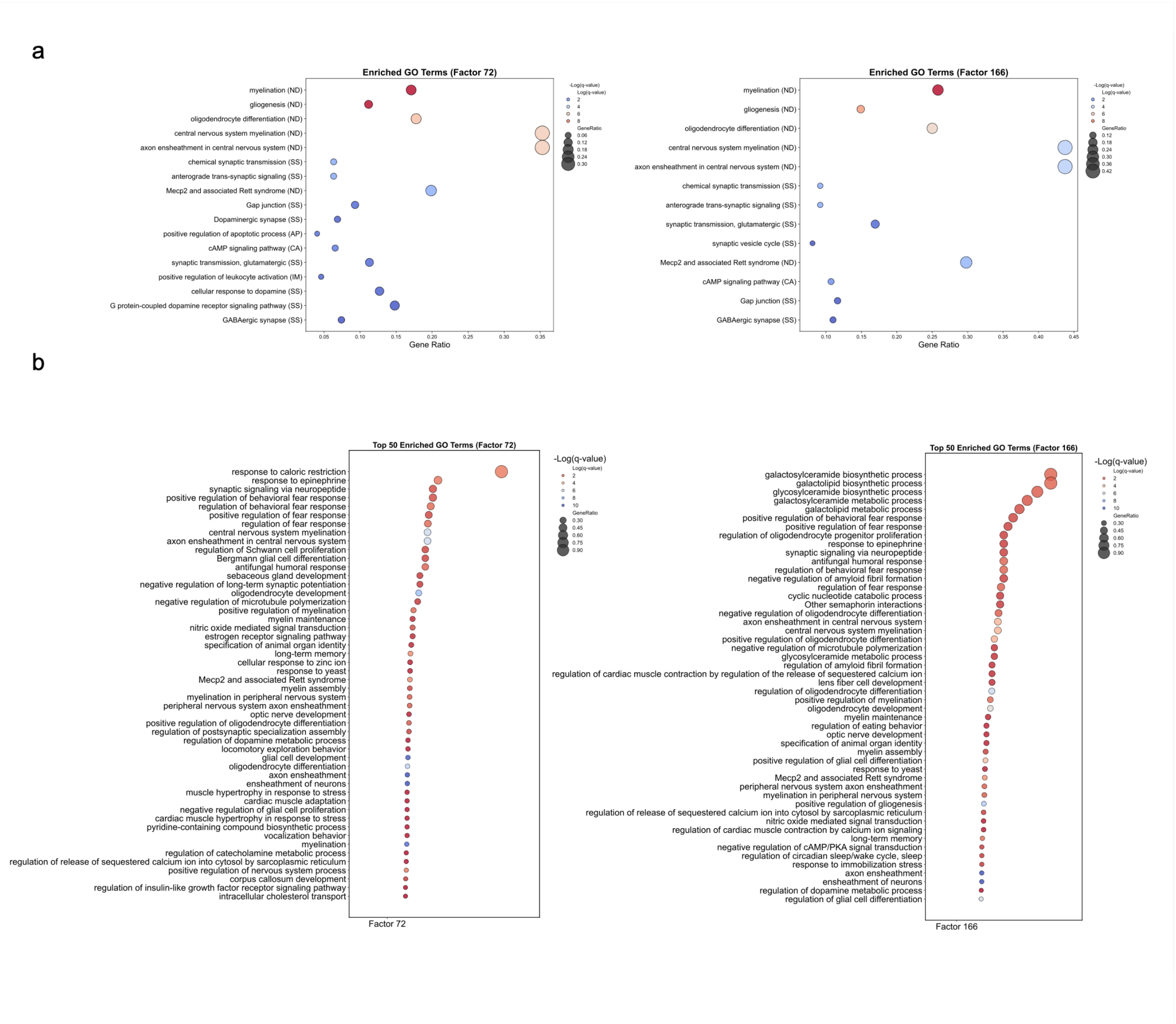


**Supplementary Fig. 4 | GO Terms for Factors 72 and 166.** **a**, Selected Go Terms; **b**, Top 50 enriched Go terms.

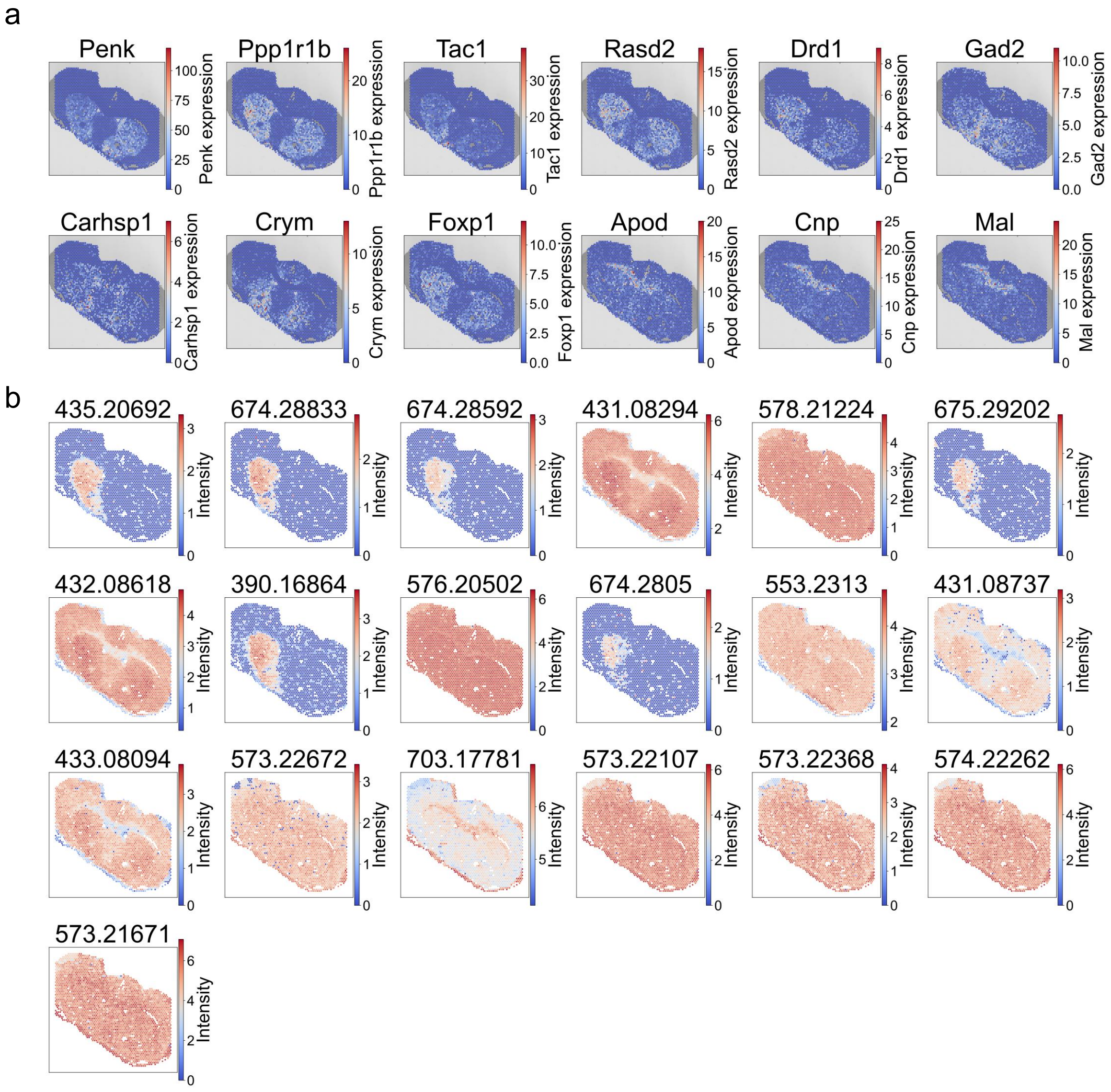


**Supplementary Fig. 5 | Spatial visualization of marker genes and metabolites of PD-related factors.** Spatial distributions of representative genes and metabolites highly associated with PD-related latent factors (e.g., Factor 72 and Factor 166), illustrating their lesion-enriched localization patterns in the striatum. **a**, Spatial expression of top-ranked genes. **b**, Spatial distribution of top-ranked metabolites (by m/z). These spatial patterns support the coordinated transcript–metabolite structure recovered by SHINE.

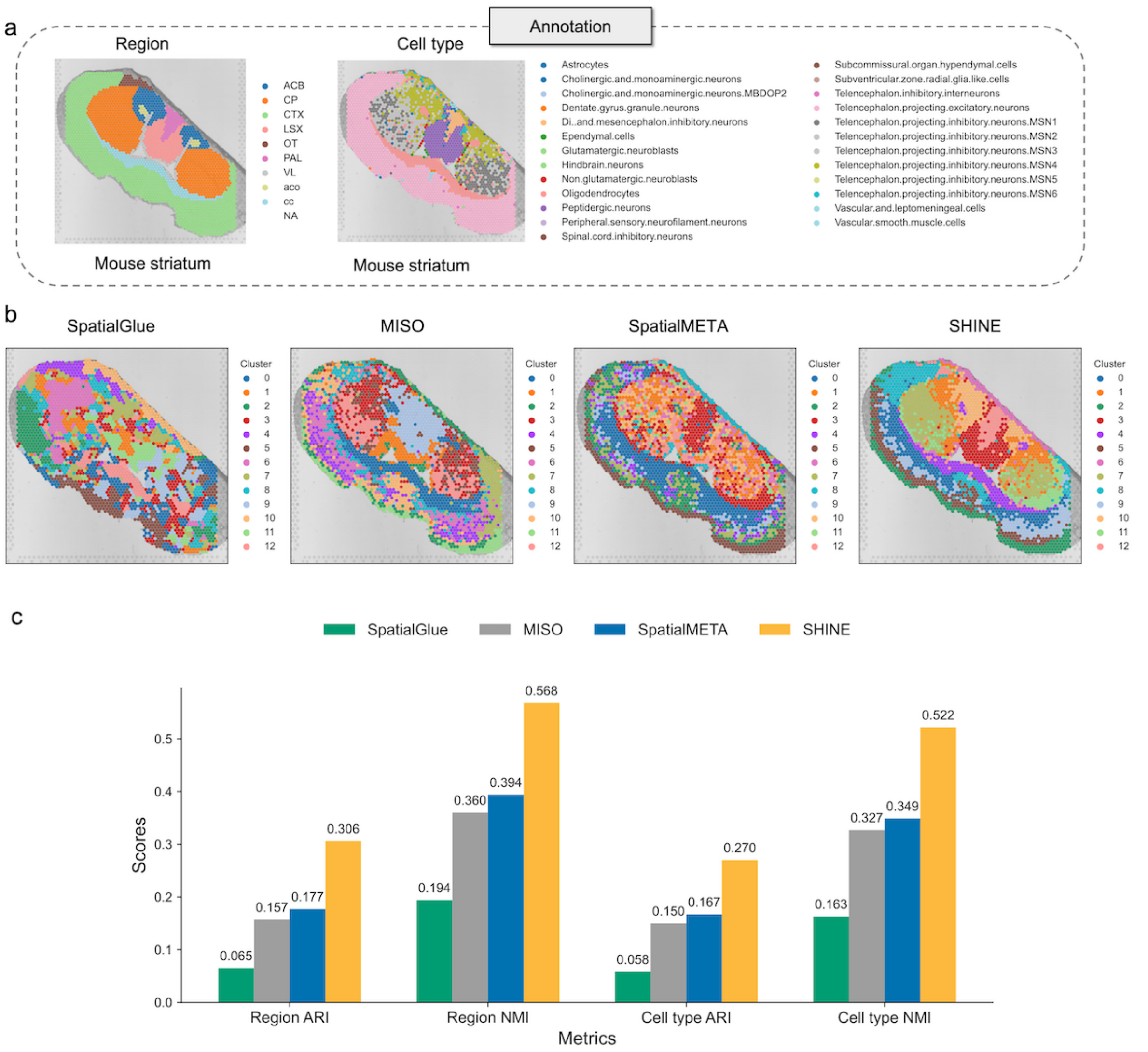


**Supplementary Fig. 6 | Comparison of integrated spatial transcriptomic and DHB matrix–assisted metabolomic clustering results with ground truth in the mouse striatum.** **a**, Spatial annotations of the mouse striatum showing anatomical regions (left) and cell-type annotations (right), used as reference labels for evaluation; **b**, Spatial clustering results generated by SpatialGlue, MISO, SpatialMETA, and SHINE on the mouse striatum tissue section; **c**, Quantitative comparison of clustering performance across methods using region- and cell-type–level agreement metrics (ARI and NMI).

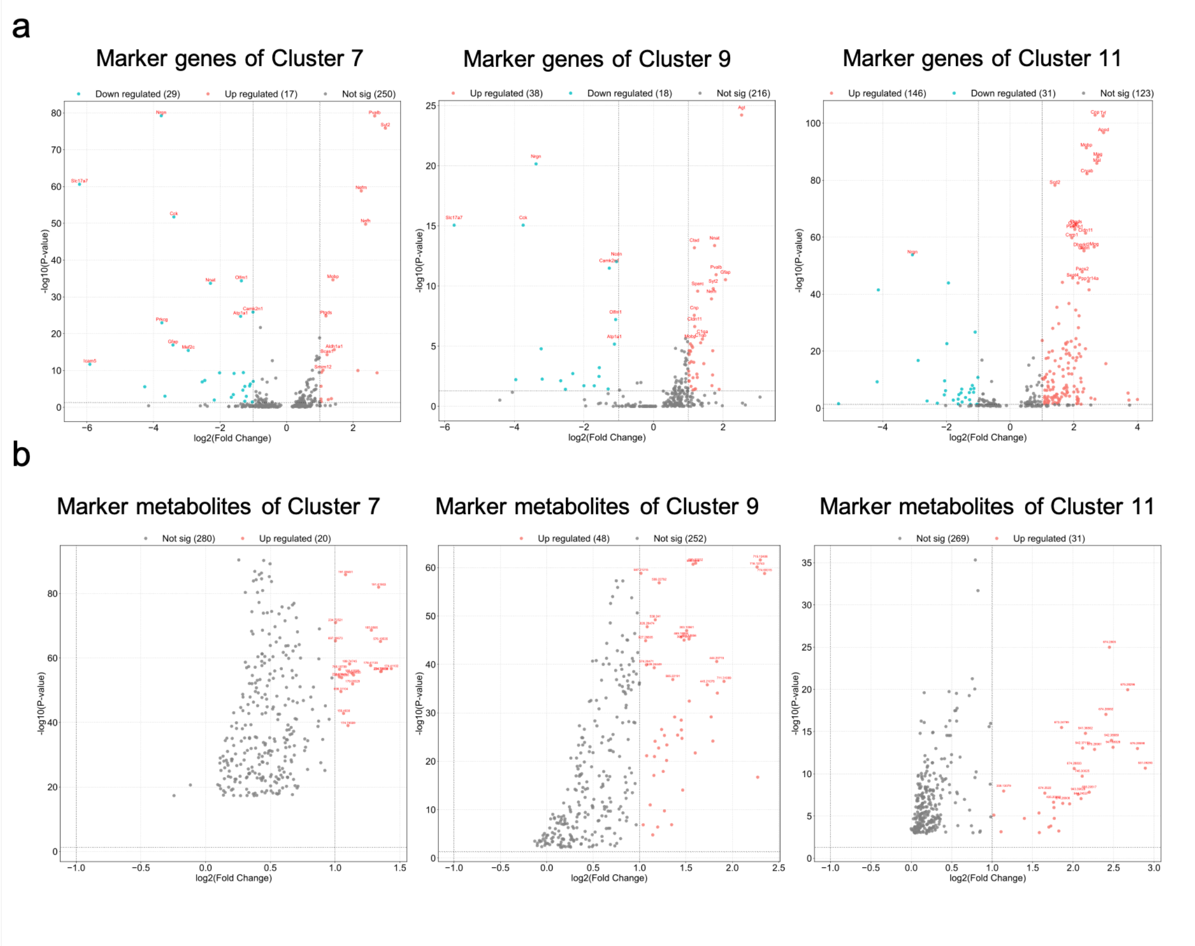


**Supplementary Fig. 7 | Differential marker genes and metabolites in PD-associated clusters.** **a**, Volcano plots showing cluster-specific marker genes for Clusters 7, 9 and 11, identified based on $\log_2$ fold change and significance thresholds. **b**, Differentially abundant metabolites corresponding to the same clusters.

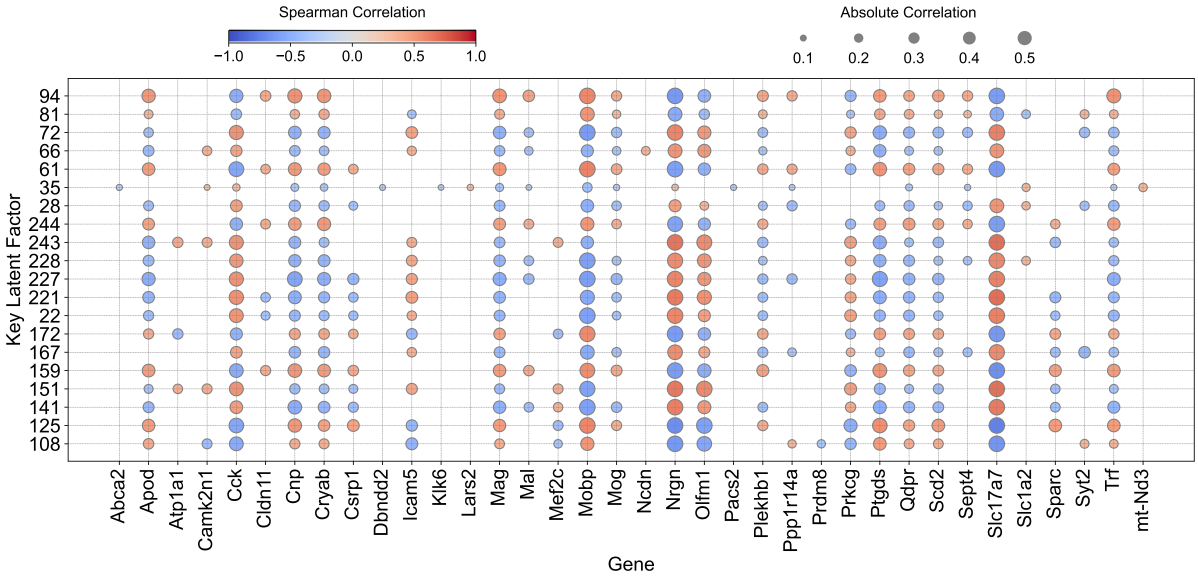


**Supplementary Fig. 8 | Latent factor–gene association landscape across substantia nigra.** Spearman correlation matrix between key latent factors (rows) and representative PD-related genes (columns). Dot size reflects absolute correlation strength; color represents directionality. Factor 94 strongly correlates with genes involved in oxidative stress, myelin regulation, and neurodegeneration (e.g., *Cryab*, *Mobp*, *Qdpr*).

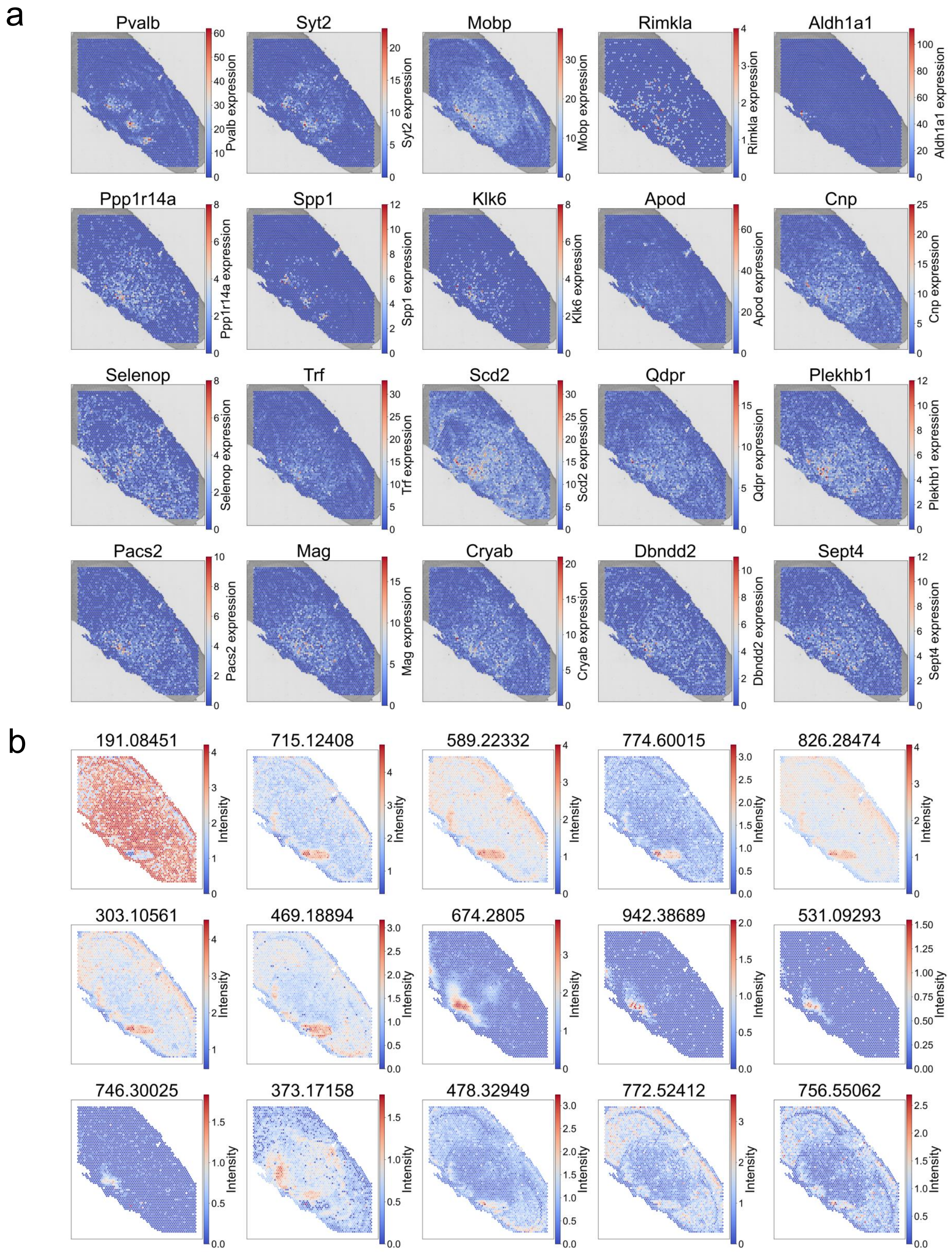


**Supplementary Fig. 9 | Spatial distribution of PD-related marker genes and metabolites.** Spatial distributions of molecular features associated with Factor 72 and Factor 166, identified based on Spearman correlation analysis, are shown. Genes and metabolites with strong associations to the latent factors were visualized to facilitate interpretation of their spatial patterns and potential Parkinson's disease relevance. **a**, Spatial expression of PD-relevant genes; **b**, Comprehensive spatial visualization of factor-associated metabolites.

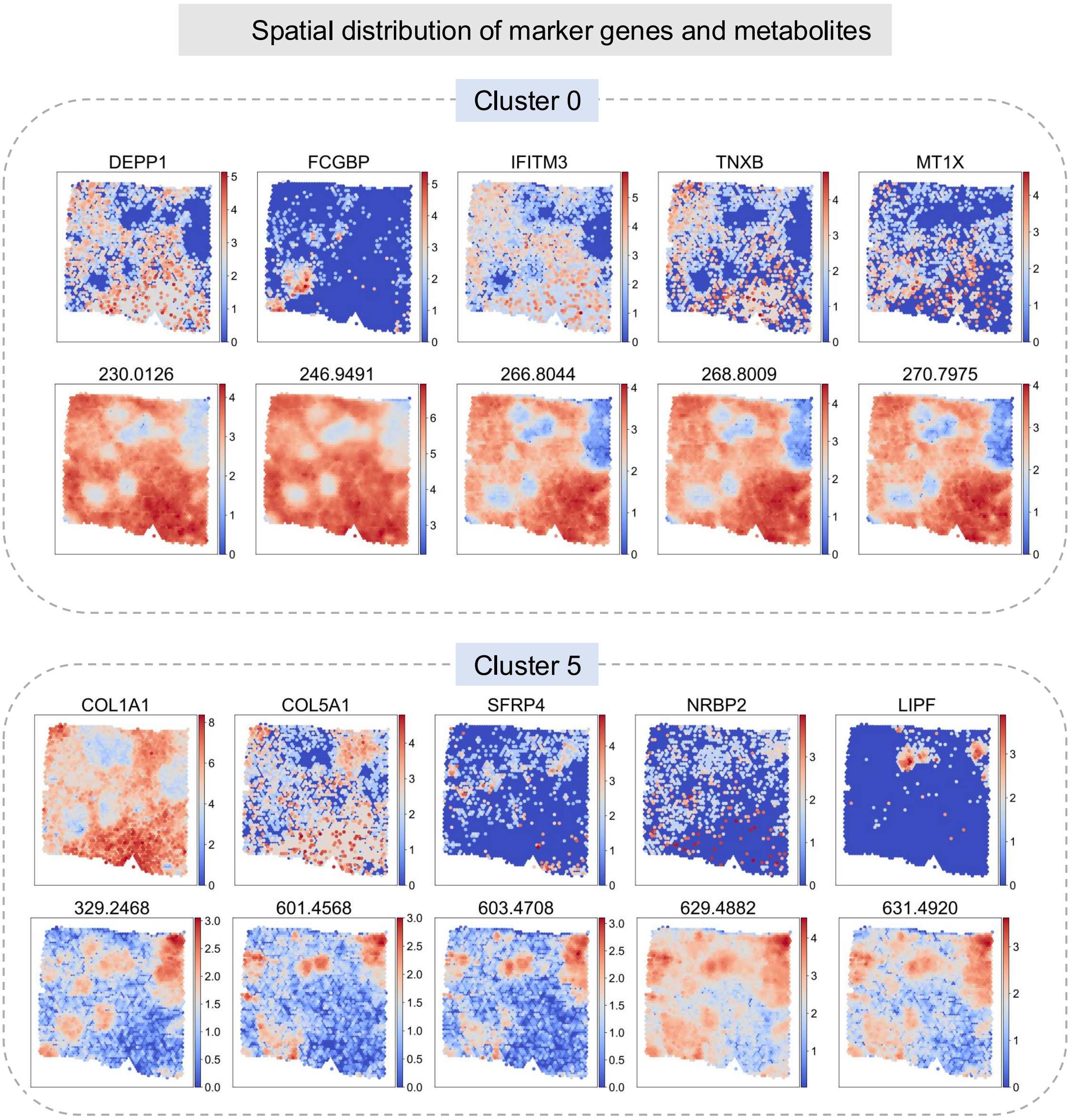


**Supplementary Fig. 10 | Spatial distribution of cluster-specific (Clusters 0&5) gene and metabolite markers identified by SHINE.**

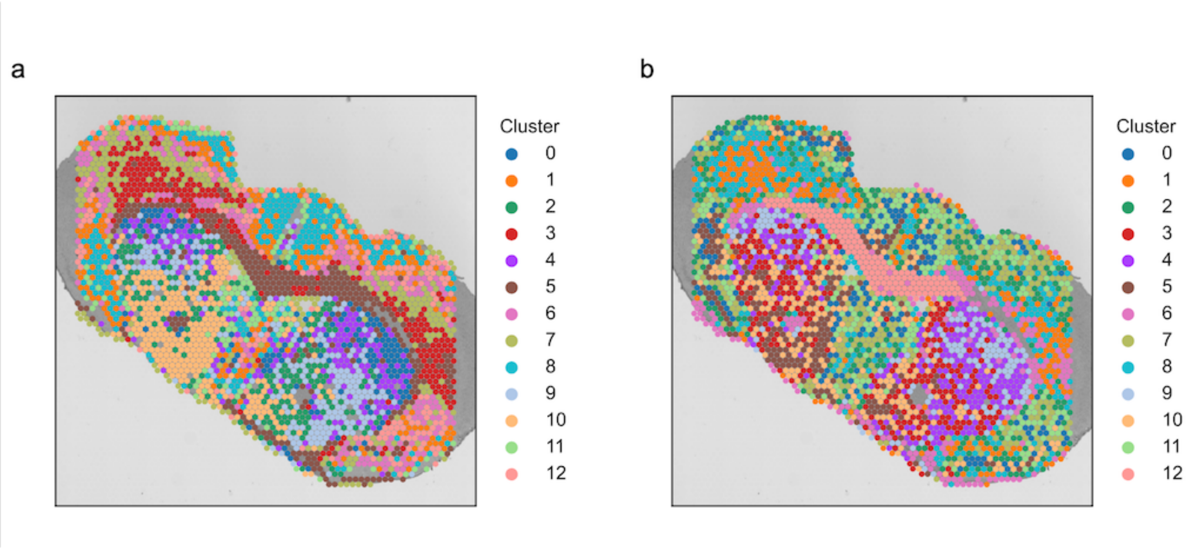


**Supplementary Fig. 11 | Spatial clustering results obtained using the SpatialMETA alignment strategy. a**, clustering results reproduced using the original implementation. **b**, clustering results obtained after performing clustering on the registered data.
